# A novel independence test for somatic alterations in cancer shows that biology drives mutual exclusivity but chance explains co-occurrence

**DOI:** 10.1101/052803

**Authors:** Sander Canisius, John W.M. Martens, Lodewyk F.A. Wessels

## Abstract

Just like recurrent somatic alterations characterize cancer genes, mutually exclusive or co-occurring alterations across genes suggest functional interactions. Identifying such patterns in large cancer studies thus helps the discovery of unknown interactions. Many studies use Fisher’s exact test or simple permutation procedures for this purpose. These tests assume identical gene alteration probabilities across tumors, which is not true for cancer. We show that violating this assumption yields many spurious co-occurrences and misses many mutual exclusivities. We present DISCOVER, a novel statistical test that addresses the limitations of existing tests. In a comparison with six published mutual exclusivity tests, DISCOVER is more sensitive while controlling its false positive rate. A pan-cancer analysis using DISCOVER finds no evidence for widespread co-occurrence. Most co-occurrences previously detected do not exceed expectation by chance. In contrast, many mutual exclusivities are identified. These cover well known genes involved in the cell cycle and growth factor signaling. Interestingly, also lesser known regulators of the cell cycle and Hedgehog signaling are identified.

**Availability:** R and Python implementations of DISCOVER, as well as Jupyter notebooks for reproducing all results and figures from this paper can be found at http://ccb.nki.nl/software/discover.

## 1 Introduction

Tumor development emerges from a gradual accumulation of somatic alterations that together enable malignant growth. As has been revealed by recent genomic profiling efforts, an immense diversity exists in the alterations that tumors acquire [1, 2]. Whether by e.g. copy number aberration, point mutation, or DNA methylation, alterations of many genes may potentially trigger transformation. Often though, the fate of a cell acquiring a certain alteration depends on other alterations already present [3]. Therefore, with an ever-expanding catalog of cancer genes, a need arises to establish how alterations in those genes interact to transform healthy cells to cancer cells. This task can be approached by statistical analyses aiming to uncover more complex, combinatorial patterns in somatic alterations.

Two such patterns are co-occurrence and mutual exclusivity. In the former, a group of genes tends to be altered simultaneously in the same tumor, whereas in the latter, mostly only one out of a group of genes is altered in a single tumor. Mutual exclusivity is frequently observed in cancer genomics data [4, 5]. Individual alterations targeting similar biological processes are believed to be mutually redundant, with one alteration being sufficient to deregulate the affected process. Identifying mutual exclusivity can therefore help in finding unknown functional interactions. With this in mind, several statistical methods have been proposed to identify significant patterns of mutual exclusivity [6–12].

Just as mutual exclusivity is interpreted as a sign of redundancy, cooccurrence is often held to entail synergy. Alteration of only one of the two genes would be relatively harmless, whereas cells with alterations in both progress to malignancy. If such synergy exists, cancer genomes should be enriched for these co-alterations, i.e. tumors harboring alterations in both genes should be more frequent than expected by chance. Several studies have reported an abundance of co-occurring somatic alterations in various types of cancer [13–18]. For somatic copy number changes, however, it has also been suggested that co-occurring alterations emerge from tumors’ overall levels of genomic disruption [19]. Indeed, tumors display a wide diversity in genomic instability, both across and within cancer types. In tumors harboring many alterations, one should not be surprised to see simultaneous alterations in any pair of genes. In contrast, two genes altered in a tumor carrying a small number of alterations might instead have resulted from a purifying selective process. Suggesting synergy as an explanation for observed co-occurrence is only reasonable if a simpler explanation like tumor-specific alteration rates can be rejected.

In this paper, we address the statistical implications of heterogeneous alteration rates across tumors for co-occurrence and mutual exclusivity detection. With extensive analyses of simulated data, we show how commonly used statistical tests are not equipped to deal with the mismatch between what is assumed by the test and what is encountered in the data. In the presence of heterogeneous alteration rates, countless spurious co-occurrences are picked up in data that are controlled not to contain any. At the same time, many instances of true mutual exclusivity are missed. Based on these observations, we introduce DISCOVER, a novel statistical independence test that incorporates the overall alteration rates of tumors to successfully solve the issues encountered with existing tests. We compared the performance of DISCOVER to that of several other published mutual exclusivity tests: *MEMo* [6], *muex* [8], *mutex* [9], *CoMEt* [10], *MEGSA* [11], and *TiMEx* [12]. Across the whole range of significance levels, DISCOVER is more sensitive while controlling the false positive rate at the specified level.

We also applied DISCOVER to a selection of more than 3,000 tumors across 12 different cancer types. Only one co-occurrence was detected that is not explained by overall rates of alteration alone. On the other hand, many more cases of mutual exclusivity were detected than would have been possible with traditional tests. The genes targeted by these alterations cover many of the core cancer pathways known to display such exclusivity. However, we also identified exclusivity among less canonical actors in the cell cycle, and among regulators of Hedgehog signaling.

## 2 Results

### 2.1 Common tests for co-occurrence or mutual exclusivity assume homogeneous alteration rates

A commonly used test for both co-occurrence and mutual exclusivity is Fisher’s exact test applied to a 2 × 2 contingency table [16–18]. The test is used to support co-occurrence when the number of tumors with alterations in both genes is significantly higher than expected by chance. Likewise, it suggests mutual exclusivity when the number of tumors with alterations in both genes is significantly lower. The validity of this test depends on the assumption that genes’ alterations across tumors are independent—a reasonable assumption— and identically distributed (i.i.d.). The latter implies that the probability of an alteration in a gene is the same for any given tumor. With cancer’s heterogeneity in mind, this assumption may prove more problematic. Surely, a gene is more likely found altered in tumors with many somatic alterations overall, than in tumors with only few such changes.

Other tests used for co-occurrence or mutual exclusivity depend on the same i.i.d. assumption as described for Fisher’s exact test. This is the case for permutation tests that estimate the expected number of tumors altered in both genes by randomly reassigning gene alterations across tumors [7, 13]. It is also true for a simple Binomial test that we will use to illustrate the consequences of violating the i.i.d. assumption. This test is depicted in Figure 1c. The alteration probability *p_i_* of a gene is estimated to be the proportion of tumors altered in that gene. For example, gene 3 in Figure 1a is altered in 2 of the 5 tumors, resulting in *p*_3_ = 0.4 (Fig. 1c). If alterations targeting two genes are independent, the probability of a tumor altered in both genes equals the product *p*_1_ · *p*_2_ of those genes’ alteration probabilities. Hence, out of *m* tumors, *m* · *p*_1_ *p*_2_ tumors are expected to harbor alterations in both genes. In the example in Figure 1a, the probability of alterations in both genes 3 and 5 would be *p*_3_ · *p*_5_ = 0.4 · 0.4 = 0.16. Therefore, if alterations of genes 3 and 5 were independent, we would expect 5 · 0.16 = 0.8 tumors with alterations in both. Observing more such tumors suggests co-occurrence, whereas observing fewer suggests mutual exclusivity (Fig. 1b).

**Figure 1:**
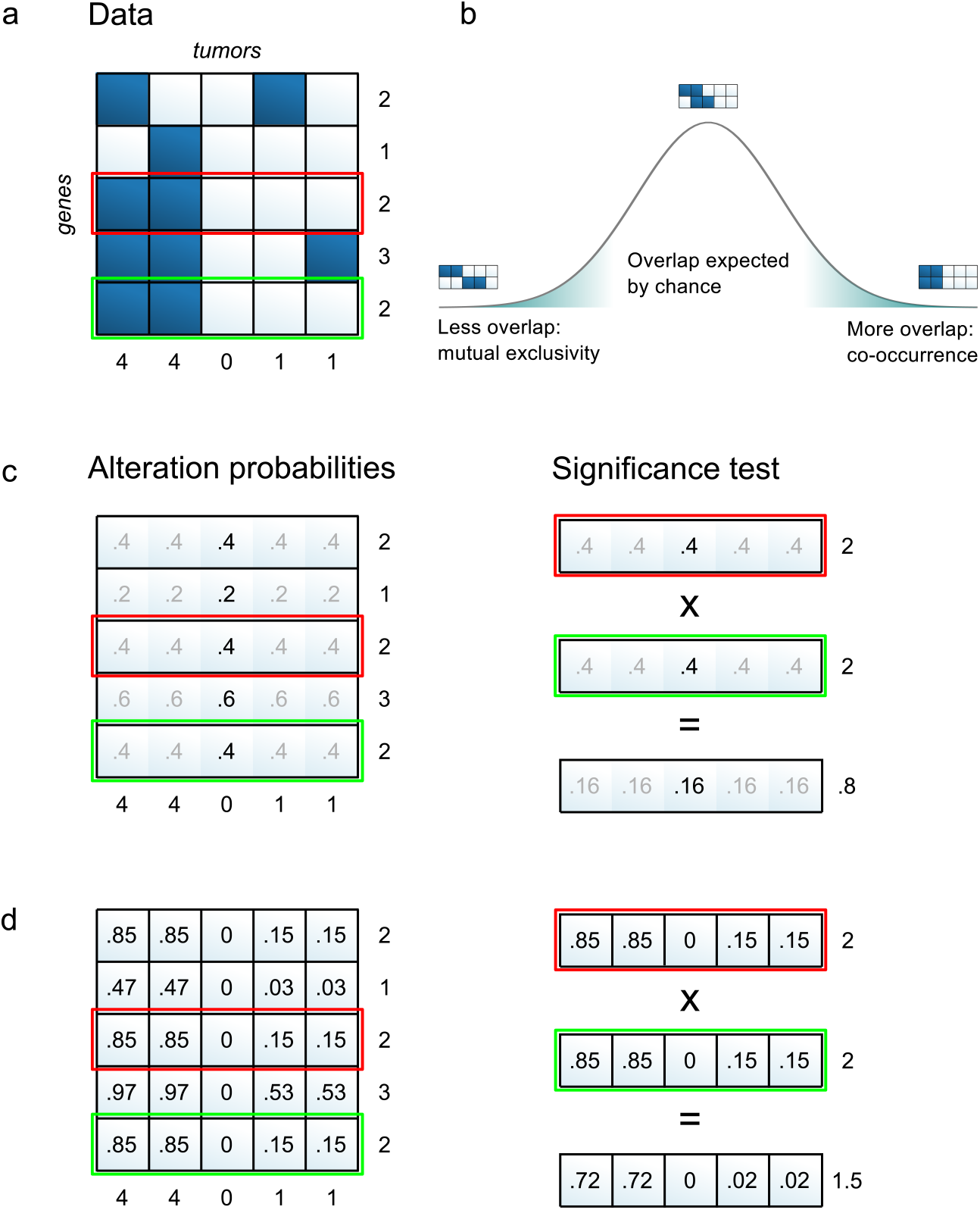
Overview of the DISCOVER method. (a) The input to the method is a binary alteration matrix with genes in the rows and tumors in the columns. The following panels illustrate how the two genes highlighted in red and green are tested for co-occurrence. (b) To identify co-occurrences or mutual exclusivities, a null distribution is estimated that describes the overlap in alterations of two genes expected by chance. Co-occurrence and mutual exclusivity correspond to the tails of this distribution. (c) In the Binomial model, a single alteration probability is estimated per gene that applies to all tumors. The expected number of alterations per gene matches the observed number. The expected number of alterations per tumor does not match the observed number. The product of two genes’ alteration probabilities gives the probability of overlap by chance, which multiplied by the number of tumors gives the expected number of tumors with alterations in both genes, in this case 0.8. (d) In the Poisson-Binomial model, gene alteration probabilities are estimated for each tumor individually. The expected number of alterations both per gene and per tumor match the observed numbers. The product of two gene alteration probabilities is also computed per tumor. The expected number of tumors with alterations in both genes according to this model is 1.5.

### 2.2 Assuming homogeneous alteration rates leads to invalid significance estimates

To illustrate the effect of the i.i.d. assumption on the detection of mutual exclusivities and co-occurrences, we performed analyses on simulated data. Genomic alterations were generated such that the alteration frequencies both per gene and per tumor resemble those observed in real tumors, but without any designed relation between the genes’ alterations—i.e. genes were simulated to be independent. As these simulated data do not contain co-occurrences or mutual exclusivities, all identified departures from independence are by definition spurious. We can therefore use these data to check the validity of the Binomial test. When testing many pairs of independently altered genes, a valid statistical test should produce *P*-values that approximately follow a uniform distribution. In contrast, when we test for co-occurrence in these data, the *P*-value distribution shows a large skew towards extremely low values (Fig. 2a). Even highly conservative significance levels will mark the majority of gene pairs as significant hits. Given that no true co-occurrences exist in the simulated data, all these hits are false positives. If we test for mutual exclusivities instead, we observe a skew towards the high end of the *P*-value spectrum (Fig. 2c).

**Figure 2:**
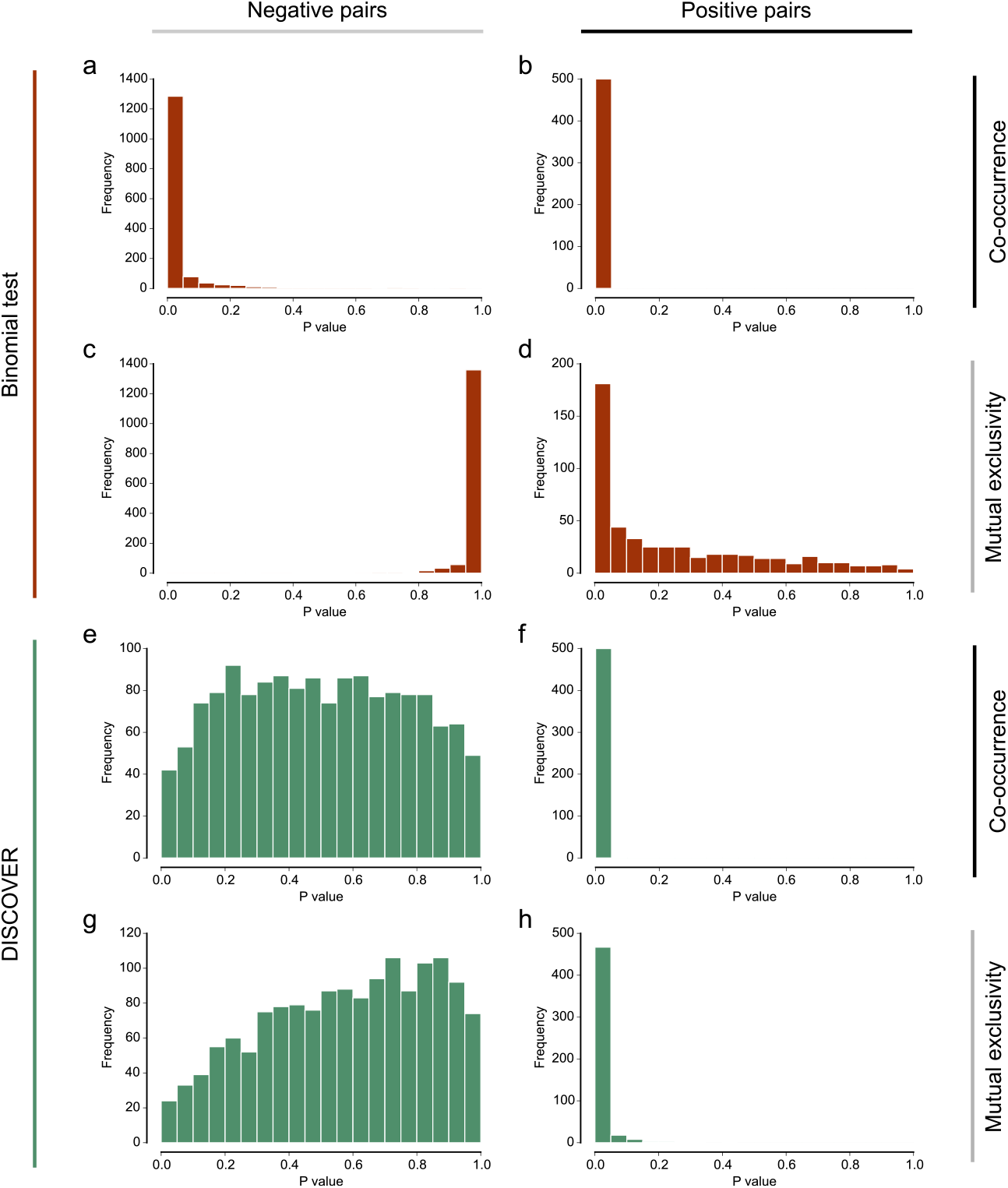
Histograms of *P*-values obtained on simulated data using either the Binomial test (a-d) or the DISCOVER test (e-h). The *P*-values apply to gene pairs with three different types of relation: gene pairs with independent alterations (a, c, e, g), gene pairs with co-occurring alterations (b, f), and gene pairs with mutually exclusive alterations (d, h).

We next evaluated the sensitivity of the Binomial test. For this, we tested simulated co-occurrences and mutual exclusivities, which we added to the data. A sensitive test should produce only low *P*-values for these positive cases, and so the resulting *P*-value distribution should be heavily skewed towards zero. If we test for co-occurrences, this is indeed the case (Fig. 2b). Testing for mutual exclusivity however, reveals a distribution that, although skewed towards lower *P*-values, is much more stretched out across the [0, 1] interval (Fig. 2d). Even highly liberal significance levels will only recover a small part of the positive cases.

We conclude that the Binomial test is anti-conservative as a co-occurrence test. In contrast, as a mutual exclusivity test, it is conservative. While we used the Binomial test for this illustration, we found the same to be true for Fisher’s exact test (Supplementary Fig. 1). To confirm our hypothesis that the i.i.d. assumption is causal to this incorrect behavior, we generated additional simulated data, making sure that the overall alteration rate is similar across the tumors. Using the Binomial test to detect co-occurrence and mutual exclusivity of independent genes results in *P*-value distributions that are much closer to uniform (Supplementary Fig. 2). This confirms that statistical tests that rely on the i.i.d. assumption are not suited for co-occurrence analysis, and have reduced sensitivity for mutual exclusivity analysis.

### 2.3 A novel statistical test for co-occurrence and mutual exclusivity

Our new method, which we call Discrete Independence Statistic Controlling for Observations with Varying Event Rates (DISCOVER), is a statistical independence test that does not assume identically distributed events. The main ingredients of the method are depicted in Figure 1d. Unlike the simpler Binomial test, we allow different tumors to have different alteration probabilities for the same gene—the alteration probabilities for genes 3 and 5 in Figure 1d now vary per tumor, in contrast to Figure 1c. For tumors with many altered genes, this probability is higher than for tumors with only few alterations. To estimate these alteration probabilities, we solve a constrained optimization problem that ensures that the probabilities are consistent with both the observed number of alterations per gene and the observed number of alterations per tumor. The probability of concurrent alterations in two independent genes is then obtained for each tumor individually, by multiplying the tumor-specific gene alteration probabilities, as indicated in the right panel of Figure 1d. With these probabilities, an analytical test based on the Poisson-Binomial distribution can be performed to decide whether the number of tumors altered in both genes deviates from the expectation.

We repeated the simulation study performed for the Binomial test, this time applying the DISCOVER test. First, our data only contained independently generated alterations. Testing for co-occurrence (Fig. 2e) and mutual exclusivity (Fig. 2g) resulted in *P*-value distributions much closer to uniform, as one would expect. The fact that these distributions are not truly uniform is a property shared by all discrete test statistics [20]; it makes discrete tests slightly more conservative. Most importantly, the anti-conservative bias towards co-occurrence of the Binomial test is not present in the DISCOVER test. By testing simulated co-occurrences, we established that the removal of the anti-conservative bias does not compromise the sensitivity for true cooccurrences (Fig. 2f). Moreover, the sensitivity for mutual exclusivities is improved when compared with the Binomial test (Fig. 2h).

### 2.4 Extension to a group-based mutual exclusivity test

Mutual exclusivity is not restricted to pairs of genes. Larger groups of genes may also display alteration patterns in which most tumors only have an alteration in one of the genes. We considered three statistics to assess the mutual exclusivity of groups of genes: coverage, exclusivity, and impurity (Fig. 3a). For all three of these statistics, its expectation for groups of independent genes can be described by a Poisson-Binomial distribution (see Methods), and thus a statistical test can be formulated for determining significance. Based on simulated data, we established that the impurity-based group test has the best balance between sensitivity and specificity (Supplementary Fig. 3).

**Figure 3:**
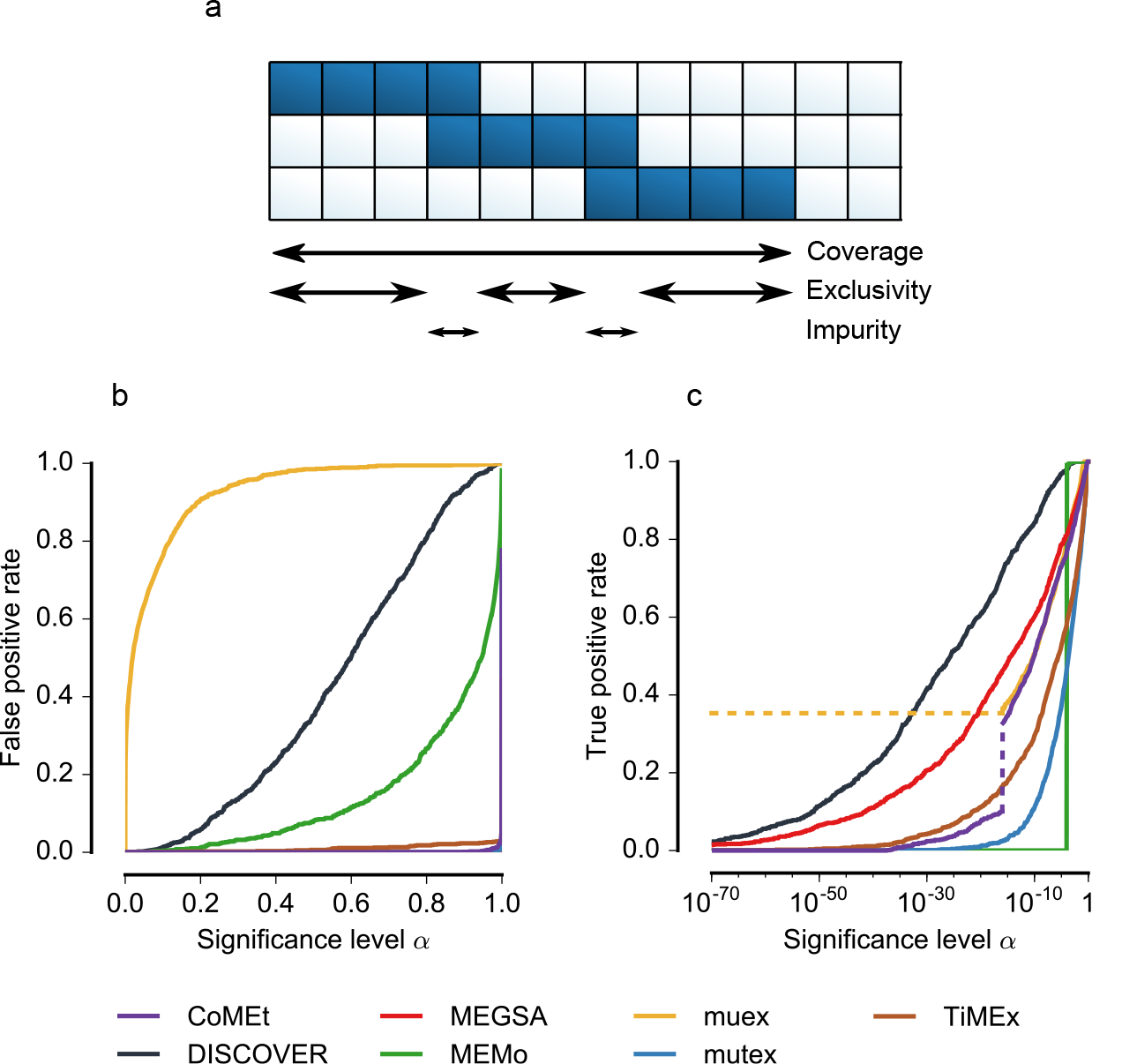
Extension of the DISCOVER test for mutual exclusivity within groups of genes. (a) Three alternative statistics for measuring the degree of mutual exclusivity within a group of genes. Coverage refers to the number of tumors that have an alteration in at least one of the genes. Exclusivity refers to the number of tumors that have an alteration in exactly one gene. Impurity refers to the number of tumors that have an alteration in more than one gene. (b) *P*-value reliability curves comparing DISCOVER with other mutual exclusivity tests. The false positive rate should not exceed the significance level *α*. In such a case, the calibration curve will be below the diagonal. For all tests but muex, this is the case. (c) Sensitivity curves comparing DISCOVER with other mutual exclusivity tests. More sensitive tests will attain higher true positive rates at lower significance levels. Two discontinuities that occur at a significance level of approximately 1 × 10^−16^ are marked with dotted lines. First, muex compresses all lower *P*-values to 0, hence all lower significance levels have the same true positive rate. Second, this significance level coincides with the change from the slower CoMEt exact test to the binomial approximation (see Methods); the two tests seem to behave quite differently.

### 2.5 Comparison to other mutual exclusivity tests

We compared the performance of the group-based DISCOVER test to that of several other published mutual exclusivity tests: *MEMo* [6], *muex* [8], *mutex* [9], *CoMEt* [10], *MEGSA* [11], and *TiMEx* [12]. In this comparison, we focused on the statistical tests for mutual exclusivity provided by these methods (see Methods). Although the tests differ in the statistical model upon which they are based, all but MEMo assume identical alteration probabilities across tumors. Like Fisher’s exact test and the binomial test, they are thus examples of tests based on the i.i.d. assumption. MEMo does take into account tumor-specific alteration rates, by preserving these rates in a permutation scheme. Unlike DISCOVER, it estimates the alteration rate with respect to a small set of recurrently altered genes as opposed to all genes.

The comparison was performed on simulated data. Groups of genes with mutually exclusive alterations of various degrees of impurity served as positive examples (see Methods). For each such group, we also selected groups of independent genes of the same size and matched to have similar alteration frequencies, to serve as negative examples. In total, 10 data sets of 100 positive and 100 negative groups were generated and evaluation metrics were averaged across these 10 sets. We evaluated the tests for both specificity and sensitivity.

To evaluate specificity, we considered the extent to which a chosen significance level *α* predicts the false positive rate obtained when groups with a nominal *P*-value less than *α* are classified as mutually exclusive. By definition of the *P*-value, rejecting the null hypothesis at a significance level *α* should guarantee that the false positive rate (or type I error rate in statistical terminology) is at most *α*. Graphically, if the false positive rate is plotted as a function of the significance level (Fig. 3b), the resulting curve would ideally follow the diagonal, or it should drop below the diagonal for more conservative tests. With the exception of muex, all methods control their false positive rate below the nominal significance level, but they do so in notably different ways. CoMEt, mutex, and TiMEx only yield false positives at extremely high significance levels. Doing so, they are more conservative than required. In contrast, DISCOVER’s curve follows the diagonal more closely. This is another confirmation that tests based on the i.i.d. assumption—like before with the binomial and Fisher’s exact tests—are more conservative than those that model the varying alteration rates. Indeed, MEMo is also less conservative than CoMEt, mutex, and TiMEx. It is more conservative than DISCOVER though, which may be explained by the different strategies for estimating the tumor-specific alteration rates: based on all genes for DISCOVER, or based on frequently altered genes only for MEMo.

To evaluate sensitivity, we compared the increase of the true positive rate as a function of the significance level (Fig. 3c). A sensitive test will already attain high true positive rates at low significance levels. Across the whole range of significance levels, DISCOVER was found to be more sensitive than any of the other tests. It identified more mutually exclusive groups at lower significance levels. Only muex initially shows a higher sensitivity, but it does so at the price of many false positives (Fig. 3b)—we suspect this is partly due to numerical imprecision. At higher significance levels, muex’s sensitivity drops below that of DISCOVER. MEMo only attains a high sensitivity at higher significance levels: it is affected by the limited resolution of its permutation test. We used 10,000 permutations, which makes the lowest possible *P*-value 1 × 10^−4^. Again contrasting tests based on their underlying assumption, we conclude that the conservatism caused by the i.i.d. assumption is reflected in a lower sensitivity. The majority of mutually exclusive groups are only identified at relatively high significance levels. If correction for multiple testing is applied, this may render many of them insignificant.

### 2.6 Co-occurrence and mutual exclusivity in pan-cancer somatic alterations

We analyzed a set of 3,386 tumors covering the 12 cancer types studied in the TCGA pan-cancer initiative [21]. An alteration matrix was constructed from recurrent copy number changes and high-confidence mutational drivers. Copy number changes were analyzed for 118 genes, of which 40 gains and 78 losses. In addition, mutation data was added for 286 genes previously classified as high-confidence driver genes [22]. In total 404 genomic alterations were analyzed covering 374 unique genes, as 30 genes are frequently targeted by both copy number changes and mutations.

We tested for pairwise co-occurrence and mutual exclusivity between pairs of genes not located on the same chromosome. These tests were stratified for cancer type to avoid confounding due to cancer type-specific alteration frequencies. Complementing the pairwise tests, we also employed the DISCOVER group test to detect patterns of mutual exclusivity in larger groups of genes. The groups we tested were selected using two different approaches. In the first approach, we extracted gene sets from the canonical pathway collection of MSigDB [23]. We tested 23 such gene sets based on pathway membership. In the second approach, we aimed to detect de novo gene sets purely based on the data. For this, we applied a clustering algorithm to the pairwise mutual exclusivity results to identify groups of genes showing a high degree of interaction.

### 2.7 No evidence for widespread co-occurrence

A remarkable outcome of our analysis is that we found no evidence for widespread co-occurrence of somatic alterations. At a maximum false discovery rate (FDR) of 1%, no significant co-occurrences were identified. Relaxing the FDR threshold to 3%, we could recover one co-occurrence, between mutation of *TP53* and amplification of *MYC*. It was recently suggested that MYC-amplified tumors show higher levels of *MYC* expression in tumors with a *TP53* mutation than in tumors without [24]. No further, reasonable relaxation of the significance threshold led to additional hits. Certainly, more gene pairs exist that harbor alterations in overlapping sets of tumors. Yet, the sizes of those overlaps do not exceed what is expected by chance if differences in tumor-specific alteration rates are taken into account. This is in sharp contrast with the significance estimates obtained with the Binomial test, which identifies 21,627 significant co-occurrences, almost one third of all pairs tested.

With the aim of establishing that the DISCOVER test is not overly conservative, we tested for co-occurrence between copy number changes of genes on the same chromosomes. Due to the inherent correlation in copy number of genes situated close to each other, such gene pairs can be considered positive controls. Indeed, all but one of the 112 pairs of tested genes located in the same recurrently altered segment are identified as co-occurring by the DISCOVER test. In addition, 18 pairs of genes situated on the same chromosome arm are detected as co-occurring, as are *DDAH1* on 1p22 and *MCL1* on 1q21. More generally, pairs within the same segment are assigned lower *P*-values on average than are pairs within the same chromosome arm (*P* = 7 × 10^−39^, Supplementary Fig. 4). The same is true, to lesser extents, for pairs within the same chromosome arm compared to pairs within the same chromosome (*P* = 6 × 10^−8^), and for pairs within the same chromosome compared to pairs across chromosomes *(P* = 0.0004).

### 2.8 Mutually exclusive alterations target core cancer pathways

Pairwise mutual exclusivities were found among 181 pairs of genes, at a maximum false discovery rate of 1% (Supplementary Table 1). We once more confirmed that detecting mutual exclusivities using the Binomial test results in far fewer significant mutual exclusivities—only three pairs were identified. Among the 181 gene pairs, there were 107 unique genes. Many of these are significantly mutually exclusive with only one or a few other genes. For some, reduced statistical power due to low alteration frequency may be the reason for not detecting more associations. However, alteration frequency is not the dominant factor in how often mutual exclusivity is detected (Fig. 4a). For example, mutations of *KRAS* are far less frequent than *TP53* or *PIK3CA* mutations. Yet, *KRAS* was found mutually exclusive with more genes than were the latter two genes.

**Figure 4:**
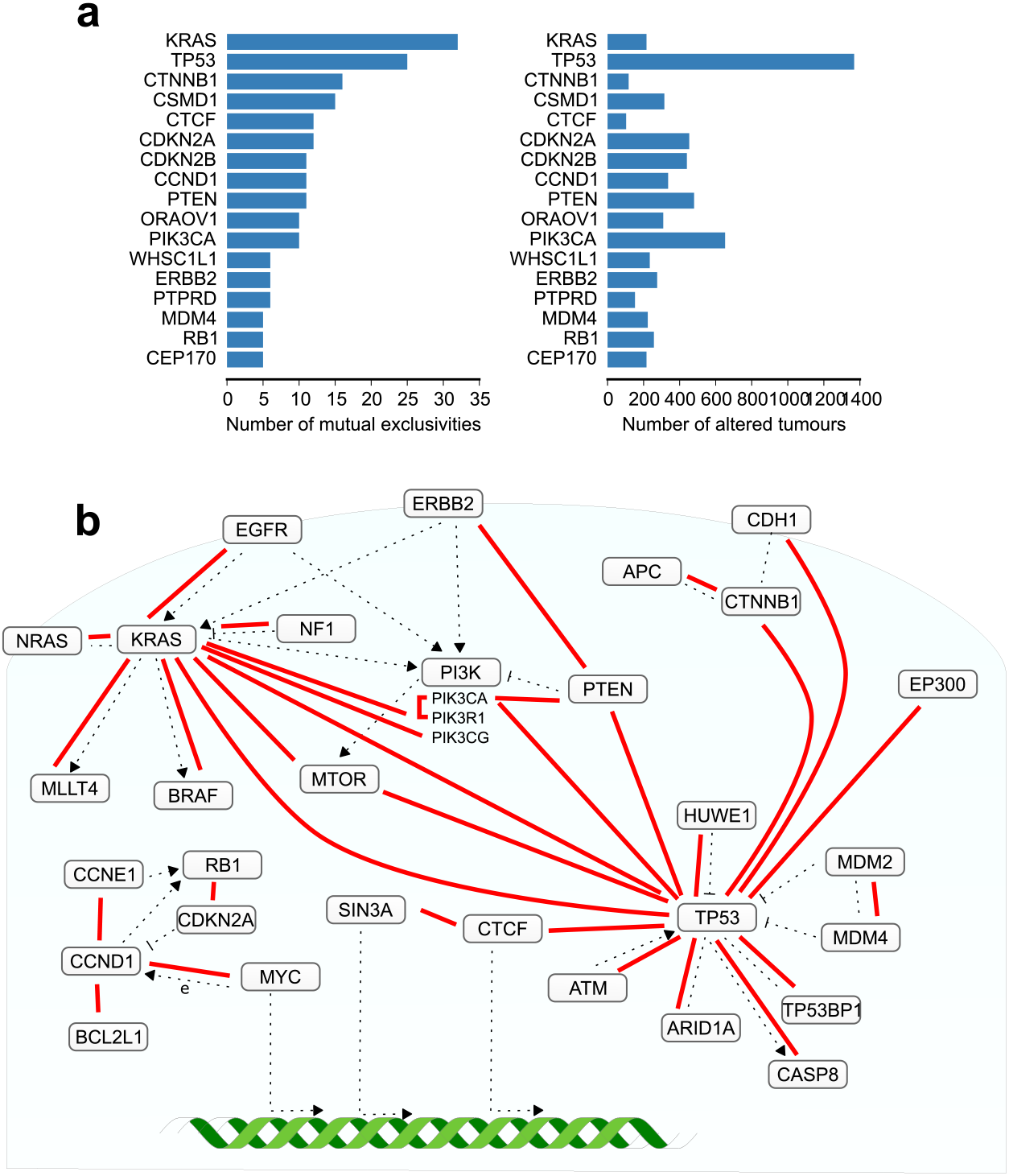
Overview of detected pairwise mutual exclusivities. (a) Comparison of the number of significant mutual exclusivities found for a gene and the number of tumors in which it has been altered. (b) Mutual exclusivities that overlap with high confidence interactions in the STRING functional interaction network depicted in their biological context. Red lines represent a mutual exclusivity between the connected genes. Dotted lines depict a functional interaction.

Since mutual exclusivity is believed often to occur between functionally related genes, we determined the overlap of the identified gene pairs with the STRING functional interaction network [25]. 31 of the identified gene pairs have a high-confidence functional interaction in STRING (Fig. 4b). This overlap is significantly higher than the 5 overlapping pairs expected by chance (*P* < 1 × 10^−4^), as determined using a permutation test. Moreover, 121 of the mutually exclusive gene pairs share a common interactor in the STRING network. By chance, this is only expected to be the case for 80 gene pairs (*P* = 0.003). This suggests that the mutual exclusivities identified are indeed for a large part driven by biological factors. The mutual exclusivities that overlap with STRING interactions revolve around three commonly deregulated processes in cancer: growth factor signaling, cell cycle control, and p53 signaling.

#### 2.8.1 Growth factor signaling

Genes coding for proteins involved in growth factor signaling are frequently altered in cancer. These alterations display a high degree of mutual exclusivity. Mutations targeting the receptor *EGFR* are mutually exclusive with mutations in its downstream mediator *KRAS*. In turn, *KRAS* mutations are mutually exclusive with mutations in its family member *NRAS*, its negative regulator *NF1*, and its downstream effector *BRAF*. All of these alterations are able to deregulate RAS signaling, and one is sufficient. Mutual exclusivity of mutations in *KRAS* and mutations in both *PIK3R1* and *PIK3CG* may be driven by the known cross-talk between RAS signaling and PI3-kinase (PI3K) signaling [26].

The PI3K signaling cascade itself is also characterized by many mutually exclusive alterations. Mutations in the *PIK3CA* and *PIK3R1* genes—both coding for components of the PI3K complex—are mutually exclusive. Alterations in the *PTEN* gene—a negative regulator of the downstream activation of AKT by PI3K—are mutually exclusive with mutations in *PIK3CA*, but also with alterations in the upstream activator of the cascade *ERBB2*. PI3K signaling is also the central biological process in several of the gene sets found mutually exclusive with the group-based test (Fig. 5a, Supplementary Fig. 5). Central genes in PI3K signaling such as *SOS1, AKT1*, and *AKT3* were not found as mutually exclusive with other pathway members in the pairwise analysis, yet the groupwise test correctly detects it.

**Figure 5:**
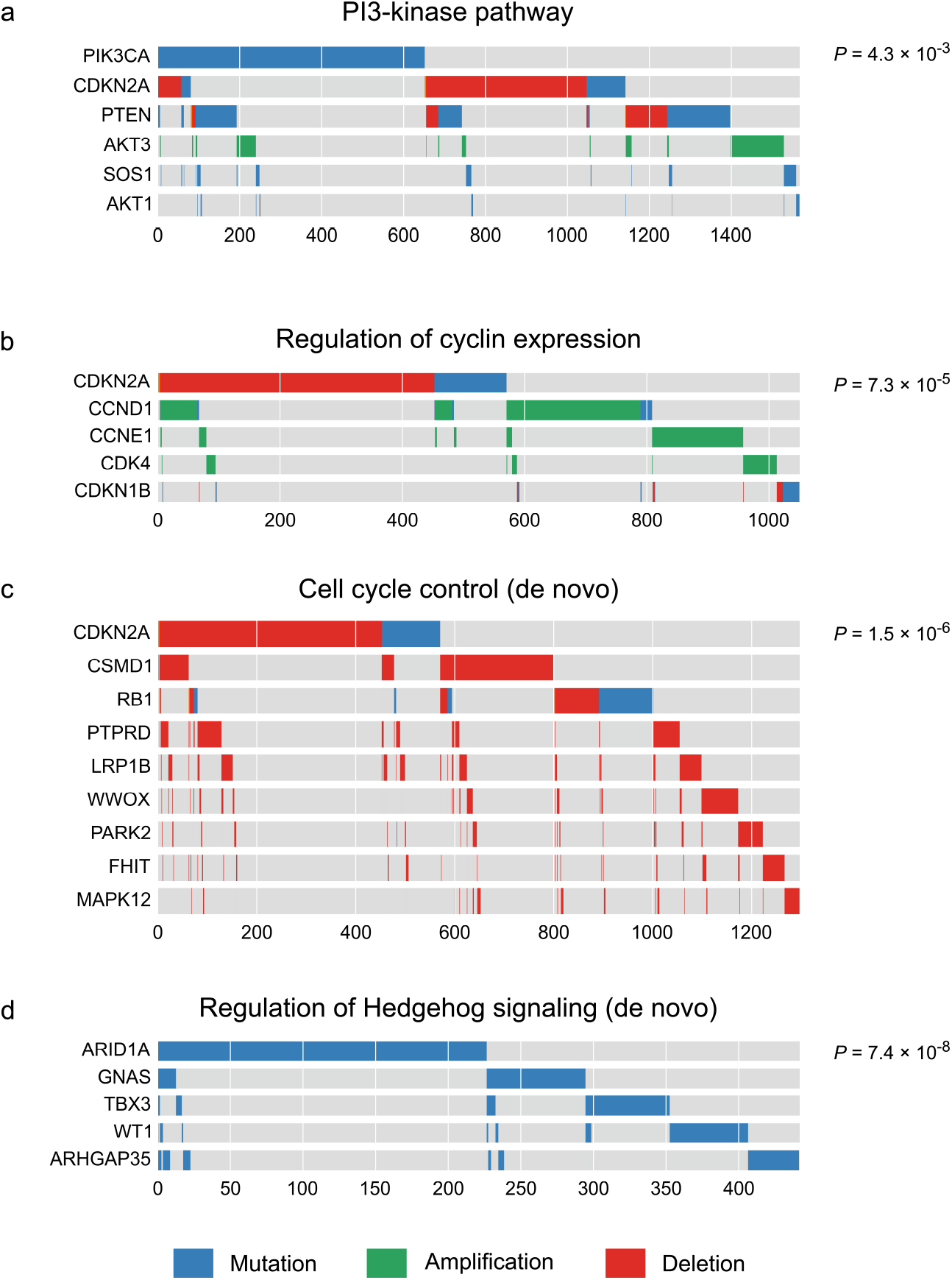
Examples of gene sets with mutually exclusive alterations. The *P*-values were computed using DISCOVER’s group-based test. Panels a and b show predefined gene sets extracted from MSigDb. Panels c and d show gene sets identified using our de novo group detection approach.

#### 2.8.2 Cell cycle control

Many tumors harbor alterations that disable the cell cycle control present in healthy cells. This control arises from a tightly regulated interplay between cell cycle activating Cyclins and CDKs, and CDK inhibitors, linked together by the master cell cycle regulator RB1. Alterations in these genes are also mutually exclusive. For example, copy number gains in *Cyclins D1* and *E1* are mutually exclusive, as are *CDKN2A* copy number loss and both mutation and copy number loss of *RB1*. The transcriptional activation of *CCND1* by *MYC* is also reflected in the mutual exclusivity between copy number gains in the two genes. Also as a group, cyclins, CDKs, and CDK inhibitors show a clear pattern of mutual exclusivity (Fig. 5b, Supplementary Fig. 5). *CDK4* and *CDKN1B*, central players in the regulation of the cell cycle, did not show up in the pairwise results, but are highly exclusive with the other genes involved.

#### 2.8.3 p53 signaling

p53 plays a pivotal role in deciding on cell fate after cellular stresses common in cancer development. For this reason, p53 mutations are the most common alterations in cancer. Not all tumors disable p53 function genetically however. Alterations in regulators of p53 provide an alternative way to deregulate p53 function in p53-wildtype tumors, but are likely redundant in tumors that already have a dysfunctional p53 protein. Indeed, we found alterations in several regulators of p53 to be mutually exclusive with *TP53* mutation. For example, mutations in its positive regulator *ATM*, but also mutations in its negative regulator *HUWE1* are mutually exclusive with *TP53* mutations. *MDM2* and *MDM4*, highly similar negative regulators of p53, have a mutually exclusive pattern of copy number gains. Mutations in *CASP8*, a downstream mediator of p53-induced apoptosis, tend also not to overlap with *TP53* mutations.

### 2.9 De novo gene set detection

As a final step in our analysis, we detected de novo gene sets purely based on observed patterns of mutual exclusivity, without input based on recorded biological knowledge. To this end, we applied correlation clustering to a network derived from pairwise mutual exclusivities (see Methods). The identified clusters were tested for groupwise mutual exclusivity with the group-based test.

One of the most significant gene sets includes *RB1* and *CDKN2A*, two pivotal players in cell cycle control (Fig. 5c). *PARK2* [27], *WWOX* [28], *FHIT* [29], *PTPRD* [30, 31], and *MAPK12* [32] have also all been linked to a regulating role in various phases of the cell cycle. They have been found to do so by regulating cyclins, CDKs, or CDK inhibitors. This functional similarity may explain these genes’ mutual exclusivity with *RB1* and *CDKN2A*. As of yet, *LRP1B* and *CSMD1* have not been linked to cell cycle control. Their mutual exclusivity with respect to several regulators of the cell cycle may instigate further study in this direction.

Another group of genes with a high degree of mutual exclusivity (*P* = 7 × 10^−8^) consists of genes that have been implicated in the regulation of Hedgehog signaling (Fig. 5d). With the exception of *ARHGAP35*, all genes in this group have experimentally been linked to a regulatory role in Hedgehog signaling. *GNAS* [33, 34], *TBX3* [35], and *WT1* [36] were found to directly regulate the pathway. *ARID1A*, coding for a component of the SWI/SNF complex, is likely to play a similar role, since loss of another component of this complex, Snf5 was found to lead to activation of the Hedgehog pathway [37]. Besides these two examples, several other gene sets were identified that combine known interaction partners with interesting leads for undiscovered interactions.

## 3 Discussion

The recent growth in the number of large genomics data sets gives rise to a parallel increase in statistical power to detect ever more complex associations. As another consequence of larger sample sizes, however, poorly matched assumptions will have an increasing impact on the results. A central assumption behind commonly used statistical tests for co-occurrence and mutual exclusivity is that a gene’s alteration probability is identical across all tumors. Using simulated data, we have shown that this assumption is not only unjustified, but that it leads to a full reversal of the associations. The Binomial test we used for illustration is but a representative of a larger class of independence tests based on the same assumption. This class includes analytical approaches such as Fisher’s exact test, CoMEt [10], and MEGSA [11], but also permutation tests where gene alterations are uniformly shuffled across the tumors.

We have presented a novel independence test based on assumptions that better match the reality of cancer genomics data. With this new test, we analyzed tumors across 12 different cancer types for the presence of co-occurrence and mutual exclusivity. Only one case of co-occurrence was found, whereas numerous cases of mutual exclusivity were detected. Performing the same analysis with the Binomial test led to the detection of many co-occurrences and almost no mutual exclusivity. Many of the mutual exclusivities missed by the Binomial test can be related to central processes in cancer biology. We found strong mutual exclusivity between genes involved in growth factor signaling and cell cycle control. Also lesser known players in the regulation of cell cycle and Hedgehog signaling were identified. Based on the results of our simulation study, we are confident that the vast majority of co-occurrences detected by the Binomial test are spurious.

The absence of widespread co-occurrence contradicts what was found in previous genome-wide studies. Besides, it seems counter to our expectation of positive selection for synergy that led us to look for co-occurrence in the first place. It is true that synergy resulting from the concurrent alteration of two genes has been observed. Concurrent mutation of two genes has been reported to act on a tumor’s response to chemotherapy, or more generally on patient survival [38, 39]. None of these phenotypes, however, have been the subject of the selection from which the original tumor emerged. Only after selective pressure for that particular phenotype has taken place—for example by treating patients—would enrichment for such co-occurrences be detected. There is no doubt cancer-driving alterations often act in concert. Yet if statistical results are to serve as support for, or even meant to identify synergy, other possible explanations for the observed co-occurrence should be accounted for. In our pan-cancer analysis, overall alteration rates explained most if not all co-occurrence.

The need to take into account higher-level structural features of samples is not unique for co-occurrence and mutual exclusivity analysis. In testing the relationship between high-dimensional gene expression data and phenotypes of interest, latent sources of heterogeneity can have a profound effect on the results. Approaches like surrogate variable analysis [40], have been developed to adjust analyses appropriately. Similarly, genome-wide association studies face the issue of latent population substructure. Again, if ignored, such substructure can drastically alter the findings. Linear mixed models have gained popularity as a method to prevent confounding [41]. Both of these examples have become standard methodologies in many biomedical analyses.

## 4 Conclusions

Co-occurrence and mutual exclusivity of somatic alterations are helpful concepts for the interpretation of cancer genomics data. For example, hypotheses about functional interactions between genes are often supported by suggested co-occurrence or mutual exclusivity of their alterations. Alarmingly, we have found that the statistical tests most commonly used for this purpose are not appropriate for testing the significance of co-occurrence. Many gene pairs believed to be co-altered more often than expected by chance, in fact do not exceed this expectation if the confounding effect of tumor-specific alteration rates is taken into account. Hypotheses formulated based on the results of those tests will therefore have limited support from the data. For this reason, we discourage the use of Fisher’s exact test or simple permutation methods for detecting co-occurrence. We have presented DISCOVER as a better alternative. Mutual exclusivity analysis using existing tests does not suffer from high false positive rates, but the sensitivity is low. DISCOVER identifies more significant mutual exclusivities without increasing the false positive rate. Thus, for both co-occurrence and mutual exclusivity analyses, we expect future cancer genomics studies to benefit from DISCOVER.

## 5 Methods

### 5.1 Independence statistic

We assess both co-occurrence and mutual exclusivity by counting how many tumors have an alteration in both genes and comparing this to the number of tumors expected to have such an overlap by chance if these alterations were independent. Importantly, the overlap expected by chance should factor in the fact that tumors with many alterations have a higher chance of such overlap than tumors with fewer alterations. Our null distribution modeling this overlap therefore takes into account both the alteration rate per gene, and the alteration rate per tumor. To this end, let *p_ij_* denote the probability of an alteration in gene *i* and tumor *j*. We assume that the alteration probability of a gene is higher in tumors with many alterations overall, than in tumors with fewer alterations. Therefore, *p_ij_* may be different from *p_ik_* for the same gene *i* in two different tumors *j* and *k*. Then, for two independent genes with alteration probabilities *p*_1*j*_ and *p*_2*j*_, the probability of an alteration in both genes in tumor *j* is *p*_1*j*_*p*_2*j*_, while for tumor *k* it is *p*_1*k*_*p*_2*k*_. Given such probabilities for a set of tumors, the number of tumors that have an alteration in both genes follows a Poisson-Binomial distribution.

The Poisson-Binomial distribution [42] describes the sum of independent, non-identically distributed Bernoulli random variables that have success probabilities *p*_1_, *p*_2_,…, *p_n_*. Its probability mass function is defined as follows.

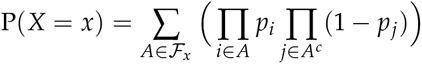

Here, 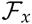 contains all subsets of size *x* of {1,2,…, *n*}, and *A^c^* denotes the complement of *A*.

Based on this distribution, we can estimate the probability of observing a number of tumors with alterations in two genes as extreme—as high for co-occurrence, or as low for mutual exclusivity—as the one observed.

If, for a given gene *i*, all probabilities *p_ij_* are equal for every tumor *j*, then the Poisson-Binomial distribution reduces to a Binomial distribution. However, estimating an individual alteration probability for every single tumor, ensures that the heterogeneity in alteration rates across tumors is taken into account.

### 5.2 Estimating gene- and tumor-specific alteration probabilities

To apply the DISCOVER test, we need estimates of the alteration probabilities *p_ij_* for all genes *i* and all tumors *j*. Let *χ* ∈ {0,1}^*n*×*m*^ denote the *n* × *m* binary alteration matrix where an entry *x_ij_* is 1 in case of an alteration in gene *i* and tumor *j*, and 0 otherwise. We use the notation *x_i•_*, and *x_•j_* for the marginal sums of the *i*th row and *j*th column respectively. Furthermore, let *X_ij_* denote the random variable for *x_ij_*, and *X_•i_*, and *X_•j_* the corresponding marginal sums. If we were to assume that the alteration of a gene is equally likely across all tumors, then the alteration probability only depends on the number of altered tumors *x_i•_*, and the total number of tumors *m*.

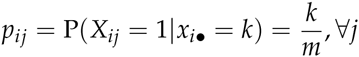

Estimating the alteration probabilities this way ensures that the expected number of alterations 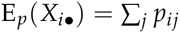 for a gene matches the observed number *x_i•_*,. In fact, the familiar expression above, is the one that maximizes the likelihood of the observed alterations under the constraint that the expected number of alterations per gene matches the observed number. To make this more explicit, we can reformulate the probability estimation as a constrained optimization problem.

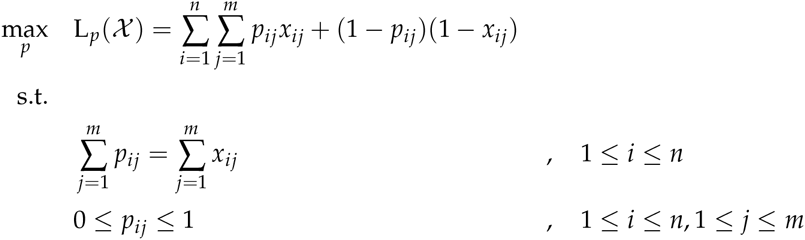

All of the above is based on the assumption that alteration probabilities for a gene are equal across tumors. Symptomatic for this assumption are probability estimates such that the expected number of alterations per tumor 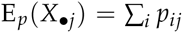 generally does not match the observed number *x_•j_*. To take into account tumor-specific alteration rates, the above optimization problem can be extended such that this expectation is also matched.

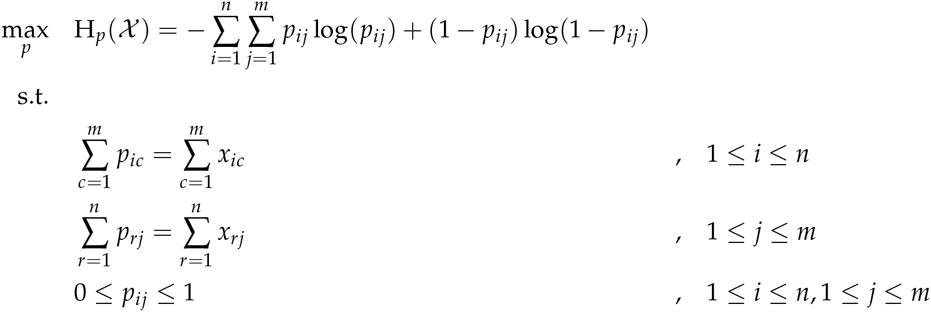

With this new formulation, the number of parameters to fit is increased by a factor *m*. As a consequence, optimizing the likelihood *L_p_(χ)* of the model risks overfitting the data. Therefore, instead of optimizing the likelihood we choose to optimize the information entropy *H_p_(χ)*. It can be shown that in the optimal solution to this reformulated problem, each alteration probability can be written in terms of two parameters (Supplementary Methods).

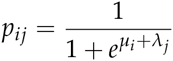

Here, each parameter *μ_i_* for gene *i* is shared by all tumors, and each parameter *λ_j_* for tumor *j* is shared by all genes. Because of this, while the original optimization problem aims to estimate *n* × *m* alteration probabilities, we can obtain the optimal solution by estimating only *n* + *m* parameters. Moreover, all genes with the same number of altered tumors share the same value for *μ_i_*. Likewise, all tumors with the same number of altered genes share the same value for *λ_j_*. This sharing of parameters leads to an even larger reduction in the effective dimensionality of the optimization.

Unlike for the Binomial case, there is no closed-form solution for estimating the *μ_i_* and *λ_j_* parameters. Instead, we use the quasi-Newton numerical optimization algorithm L-BFGS [43].

### 5.3 Stratified analysis

When the data consist of clearly separate groups of tumors, such as is the case in the pan-cancer analyses with its different cancer types, it is preferable to stratify the analysis on these groups. If for example in the mutual exclusivity analysis, group structure is not taken into account, the detected mutual exclusivities may be little more than markers for the underlying cancer types, rather than biologically related genes. The DISCOVER test is easily stratified for different groups by solving the constrained optimization problem separately for the tumors of each group. The group-specific background matrices can then be concatenated to construct a single global, but stratified, parameter matrix.

### 5.4 False discovery rate control

Commonly used procedures for multiple testing correction assume that the *P*-values are distributed uniformly under the null hypothesis. This is the case for e.g. Bonferroni correction and the Benjamini-Hochberg procedure. However, hypothesis tests that are based on a discrete test statistic, such as our DISCOVER test, are known to lead to non-uniform *P*-value distributions under the null hypothesis. In fact, pooling the *P*-values across tests with a large set of different parameters results in a *P*-value distribution that is skewed towards 1.0. This complicates the application of the standard procedures for multiple testing correction. While these procedures would still control the family-wise error rate or false discovery rate at the specified threshold, they will be more conservative because of the non-uniformity caused by the discrete test statistic. For the analyses in this paper, we used an adaptation of the Benjamini-Hochberg procedure for discrete test statistics [44].

### 5.5 Group-based mutual exclusivity test

We have defined a family of group-based mutual exclusivity tests. The following statistics can be used to assess groupwise mutual exclusivity. Each of these statistics can be shown to follow a Poisson-Binomial distribution, which we make use of to estimate significance.

- *Coverage:* the number of tumors that have an alteration in at least one of the genes. Significance is based on the probability of observing a coverage at least as high in independent genes. The Poisson-Binomial parameters for a group of genes {*g_i_* | *i* £ *I*} can be derived from the individual gene alteration probabilities as follows.

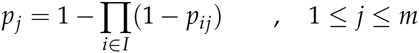 That is, the probably of at least one alteration is one minus the probability of not having any alteration.
- *Exclusivity*: the number of tumors that have an alteration in exactly one of the genes. Significance is based on the probability of observing exclusivity at least as high in independent genes. The Poisson-Binomial parameters can be derived from the gene alteration probabilities as follows.

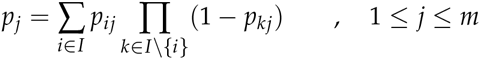
- *Impurity*: the number of tumors that have an alteration in more than one gene. Significance is based on the probability of observing impurity at least as low in independent genes. The Poisson-Binomial parameters can be derived from the gene alteration probabilities as follows.

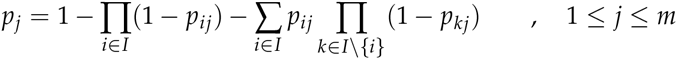 That is, the probability of more than one alteration is one minus the probabilities of no alterations and exactly one alteration. As a special case of this, if a group of only two genes is tested, the above expression reduces to *p_j_* = *p*_1*j*_ *p*_2*j*_. This is the same parameterization as was used for the pairwise test.

### 5.6 Simulation data

An alteration matrix was constructed such that alteration frequencies across both genes and tumors resembled those of real tumors. For this, we used the copy number data of the TCGA breast cancer study as a reference. Based on the copy number matrix for 24,174 genes and 1,044 tumors, we constructed two sequences of marginal counts corresponding to the number of amplifications across genes and across tumors. These two sequences were used as degree sequences to construct a random bipartite graph following the configuration model. The adjacency matrix of this bipartite graph was then used as alteration matrix for the simulated data analyses. Because of the way this matrix was constructed, the alteration frequencies across both genes and tumors resemble those of the breast cancer tumors used for reference, yet there is no dependence between alterations across genes. For the analyses, only genes with at least 50 alterations were tested.

Mutually exclusive and co-occurring gene pairs, as well as mutually exclusive gene sets were generated based on two parameters: coverage, the number of tumors altered in at least one of the genes; and impurity or overlap, the proportion of covered tumors altered in more than one of the genes. To generate pairs of mutually exclusive genes, we used quantile regression to relate the coverage of independent gene pairs to their to impurity. Simulated mutually exclusive gene pairs were generated such that their impurity was below the first percentile predicted by the quantile regression model based on their coverage. Likewise, pairs of co-occurring genes were generated such that the number of tumors altered in both genes exceeded the 99th percentile based on the coverage of independent gene pairs.

Mutually exclusive gene sets were generated by first constructing sets of purely mutually exclusive gene alterations and then adding additional, nonexclusive alterations to obtain a prespecified degree of impurity. For the former, the percentage of covered tumors was randomly sampled from a truncated normal distribution with mean 0.4 and standard deviation 0.2, truncated on the interval [0.2, 0.8]. Next, individual gene alteration frequencies were sampled from the empirical distribution of alteration frequencies in the TCGA breast cancer matrix. Gene alteration frequencies were sampled until their sum reached the coverage of the group. The number of genes thus depends on the coverage in a way that is based on realistic cancer data. As some of the mutual exclusivity tests we compared with become intractable with larger numbers of genes, we restricted the maximum number of genes to 6. In addition, we also used a minimum gene set size of 3. Finally, the impurity was sampled from the set {0.02,0.05,0.08}. Impure alterations, i.e. additional alterations in an already covered tumor, were assigned to tumors with a probability proportional to the tumor’s overall alteration frequency.

For all analyses, the background matrix for the DISCOVER test was estimated on the complete alteration matrix, including genes with fewer than 50 alterations, and including simulated co-occurrences or mutual exclusivities.

### 5.7 Comparison to other mutual exclusivity tests

We compared the performance of the group-based DISCOVER test to that of MEMo [6], muex [8], mutex [9], CoMEt [10], MEGSA [11], and TiMEx [12]. Some of these methods do more than just testing for mutual exclusivity. They combine a statistical test for mutual exclusivity with an algorithm that identifies groups of genes to test. In our comparison, we were interested in comparing the performance of the statistical tests only. We therefore evaluated the mutual exclusivity tests by applying them to pre-identified groups of genes.

For muex, MEGSA, and TiMEx, we used the R implementations provided with their respective publications. For CoMEt, we used a modified version of the official software implementation. Due to the computational complexity of the CoMEt test, it became intractable for some of the gene sets in the comparison. For this reason, the CoMEt publication suggests a set of heuristics to decide between the exact test and a faster binomial approximation, but we found those to be inadequate in our comparison. Instead, we changed the implementation such that it interrupts the CoMEt exact test after 1 minute and returns the *P*-value obtained with the binomial approximation. For the MEMo and mutex tests, we used our own implementations, which we verified to give the same results as their original Java implementations.

### 5.8 Pan-cancer alteration data

Preprocessed somatic mutation and copy number data for the 12 cancer types studied in the TCGA pan-cancer initiative [21] were obtained via Firehose (analysis run 2014_07_15). Mutations were extracted from the input of the MutSig 2CV analysis. Mutations for genes that have previously been identified as high-confidence mutational drivers [22] were included in the analysis. Discretized copy number changes were extracted from the output of GISTIC2. We considered genes altered if GISTIC2 qualified their copy number change as high-level. Pan-cancer recurrently altered regions were obtained via Synapse (syn2203662). For each region, we selected their most likely driver genes for inclusion in the analysis. If a region contained only one gene, this gene was assumed its driver. In the case of more genes, genes were selected if they overlapped with the list of high-confidence mutational driver genes, or with a curated list of cancer genes (http://www.bushmanlab.org/links/genelists).

Background matrices for the DISCOVER test were estimated for each type of alteration—mutation, amplification, and deletion—separately, and based on the genome-wide alteration matrices before gene selection. Stratification for the 12 different cancer types was applied as described before. The background matrix used in the analysis was subsequently composed out of the relevant rows in the three alteration type-specific background matrices.

### 5.9 Overlap with the STRING functional interaction network

Version 10.0 of the STRING network [25] was used to determine overlap of detected mutual exclusivities and functional interactions. We constructed a functional interaction graph by connecting genes with an edge if they have a high-confidence STRING interaction, defined by a combined score greater than 800. A mutual exclusivity graph was constructed by connecting genes with an edge if alterations in these genes were found mutually exclusive at a maximum FDR of 1%. The overlap corresponds to the number of edges appearing in both graphs. To determine enrichment of this overlap, we estimated a null distribution by randomly shuffling the gene labels of the mutual exclusivity graph 10,000 times and computing the overlap of these shuffled mutual exclusivity graphs with the unshuffled functional interaction graph.

### 5.10 De novo gene set detection

Our algorithm for detecting de novo sets of mutually exclusive genes combines two ideas from community detection. Its goal is to detect gene sets with a high likelihood of being mutually exclusive based on the results of a pairwise mutual exclusivity analysis. There are three main steps. First, a mutual exclusivity graph is constructed where genes are connected by an edge if their alterations have been identified as mutually exclusive by the pairwise test. For this step, we used a permissive significance criterion—a maximum FDR of 10%—so as not to exclude potentially interesting gene pairs that may simply not have reached significance due to the limited sample size. Second, groups of genes with a high density of mutual exclusivity edges between them are identified using a graph partitioning algorithm. Finally, these groups are subjected to the groupwise mutual exclusivity test to retain only those groups that are mutually exclusive as a group.

The graph partitioning step is based on overlapping correlation clustering. In correlation clustering, nodes in a graph are clustered such that the combined weight of edges within clusters is maximized, and the combined weight of edges between clusters is minimized. The particular algorithm we used [45] allows nodes to be assigned to multiple clusters. Moreover, we modified the original algorithm such that groups of nodes can be designated that should always share the same cluster assignments. We used this for two situations. First, genes in the same copy number segment have highly correlated copy number alterations, and consequently highly similar mutual exclusivities. Purely based on genomic data there is no reason to prefer one gene over the other, which is why we always assign all such genes to the same cluster. Second, we assume copy number alterations and mutations targeting the same gene serve the same function, and therefore add the constraint that these are always assigned to the same cluster.

The edge weights of the mutual exclusivity graph play an important role in the objective function of correlation clustering. A common phenomenon in pairwise associations is that one gene is found mutually exclusive with many other genes, but those genes are not all mutually exclusive with each other. The edges connecting the former gene may therefore not be indicative of gene set membership. They should be assigned a lower weight than edges that more specifically connect genes with a high degree of internal connectivity. To this aim, we selected the edge weights to optimize a modularity objective. In modularity optimization, a graph is compared with random graphs having the same number of nodes, edges, and degree distribution. Edges that are specific to the graph being partitioned are preferably kept within clusters, whereas edges that also appear in many of the random graphs will often span two clusters. We used a modularity measure based on conditional expected models [46]. This measure ensures that edges connecting sets of nodes with high node degrees receive a lower weight than edges that connect sets of nodes with low node degrees. In addition, it also allows for the covariance between the mutual exclusivity tests to be taken into account.

## Supplementary figures

**Supplementary Figure 1:**
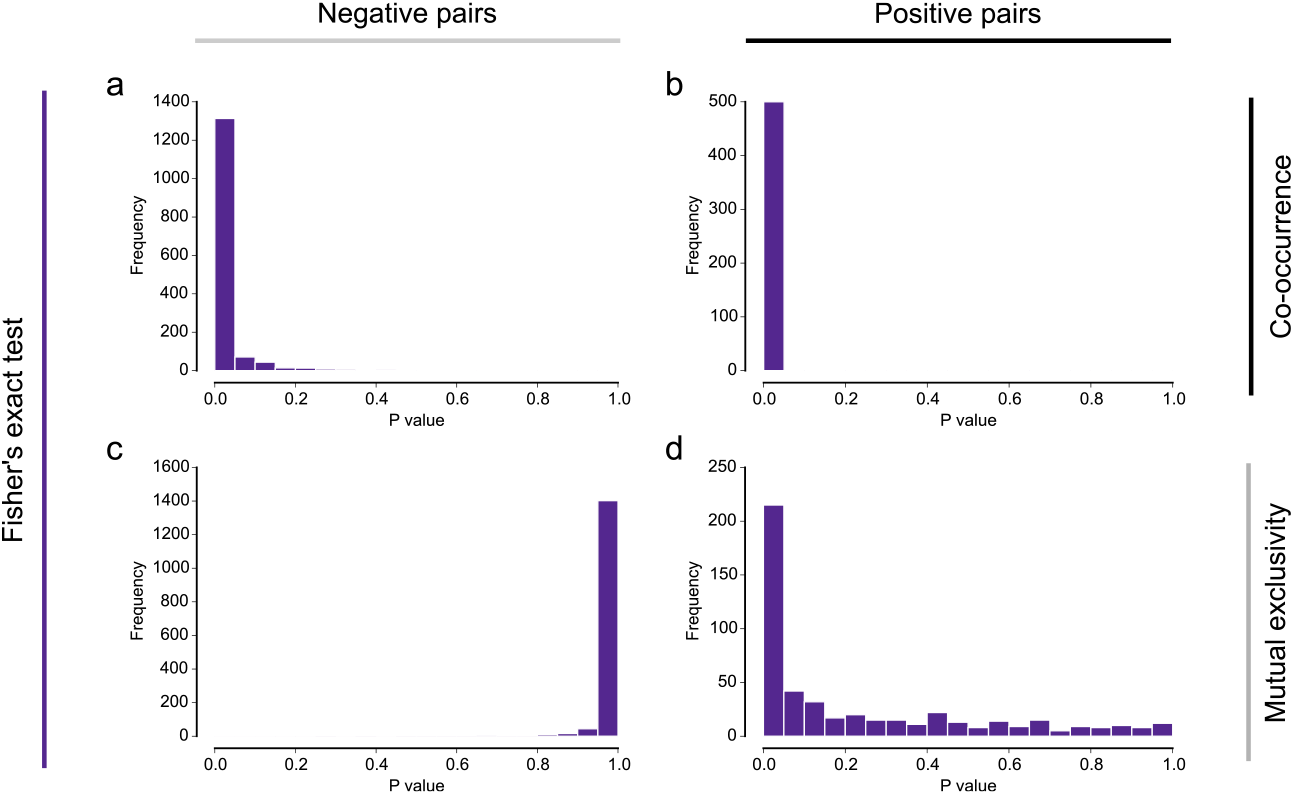
Histograms of *P*-values obtained on simulated data using Fisher’s exact test. The *P*-values apply to gene pairs with three different types of relation: gene pairs with independent alterations (a, c), gene pairs with co-occurring alterations (b), and gene pairs with mutually exclusive alterations (d).

**Supplementary Figure 2:**
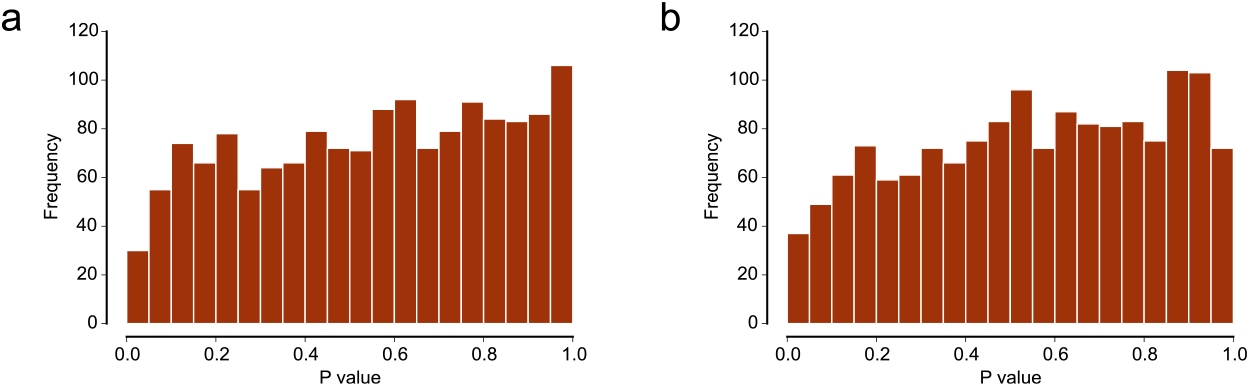
Histograms of *P*-values obtained by testing independent gene pairs for either co-occurrence (a) or mutual exclusivity (b) using the Binomial test. Simulated alteration data were generated in such a way that gene alteration frequencies resemble those in real tumors. Alteration frequencies of tumors are similar for all tumors.

**Supplementary Figure 3:**
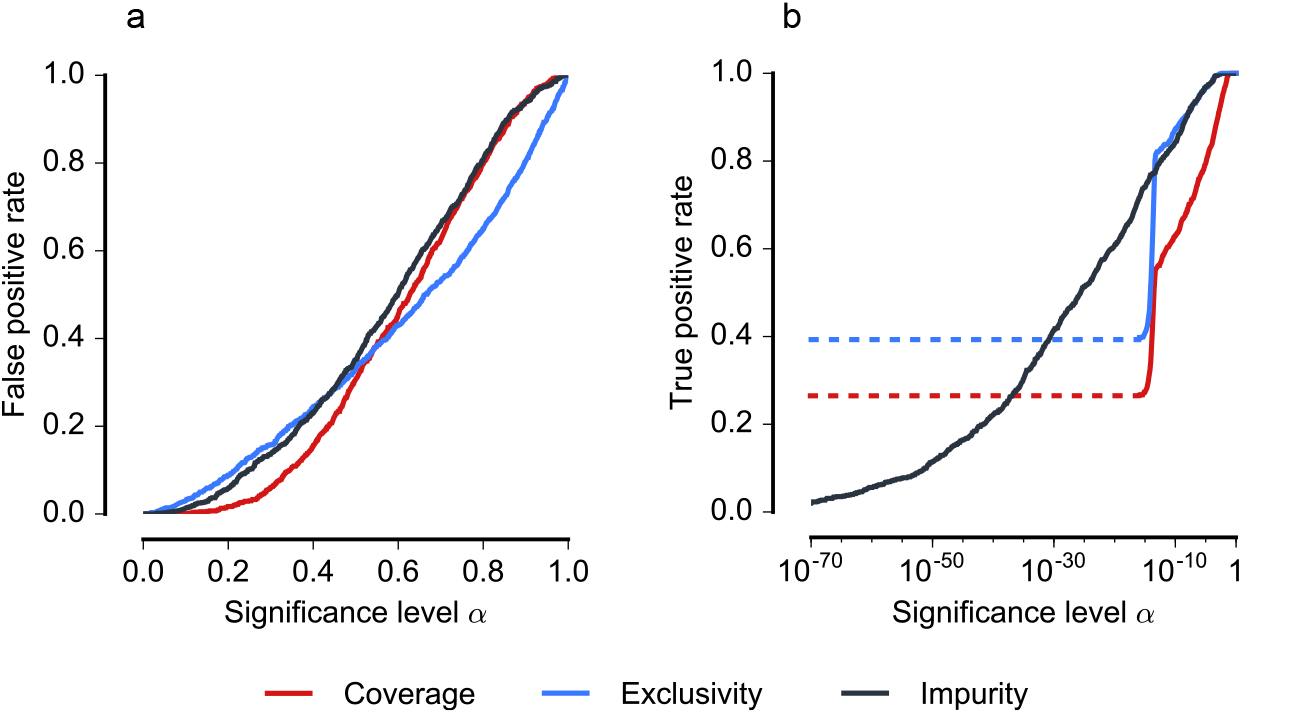
Performance on simulated gene sets of the DISCOVER test based on three alternative statistics: Coverage, the number of tumors that have an alteration in at least one of the genes; Exclusivity, the number of tumors that have an alteration in exactly one gene; Impurity, the number of tumors that have an alteration in more than one gene. (a) *P*-value reliability curves. The false positive rate should not exceed the significance level *α*. In such a case, the calibration curve will be below the diagonal. This is the case for all three statistics. (b) Sensitivity curves: more sensitive tests will attain higher true positive rates at lower significance levels. The dotted lines mark a range of significance levels where all lower *P*-values are 0.

**Supplementary Figure 4:**
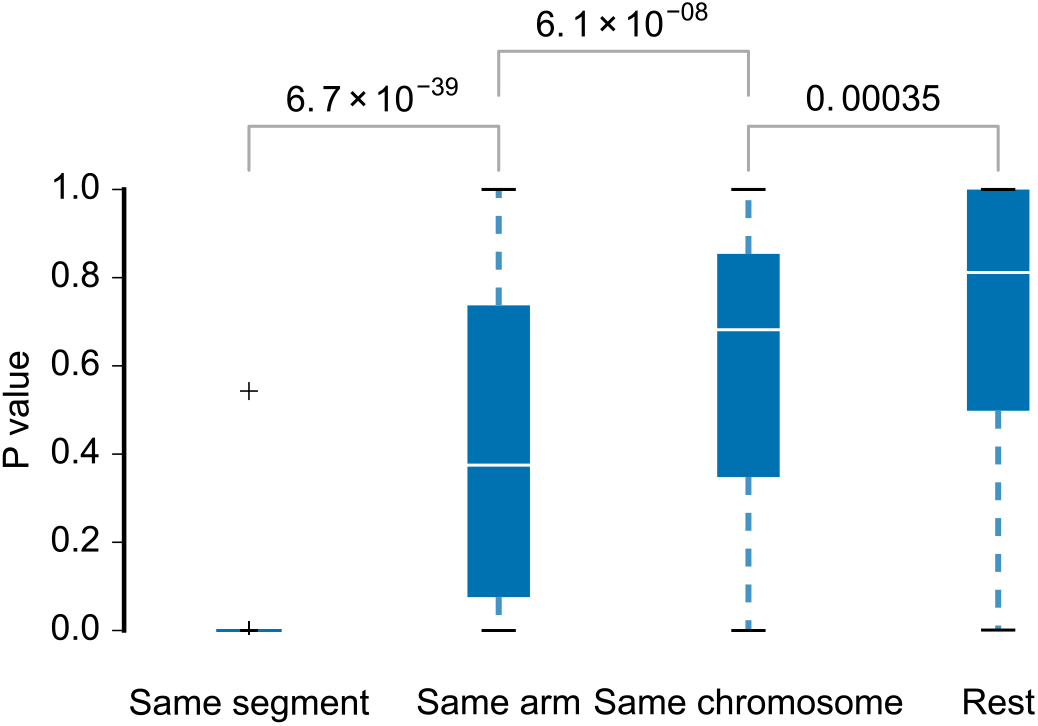
Comparison of *P*-values obtained when testing for co-occurrence between genes within the same recurrently altered segment, within the same chromosome arm, within the same chromosome, and across chromosomes. Differences in mean are tested with the Mann-Whitney *U* test.

**Supplementary Figure 5:**
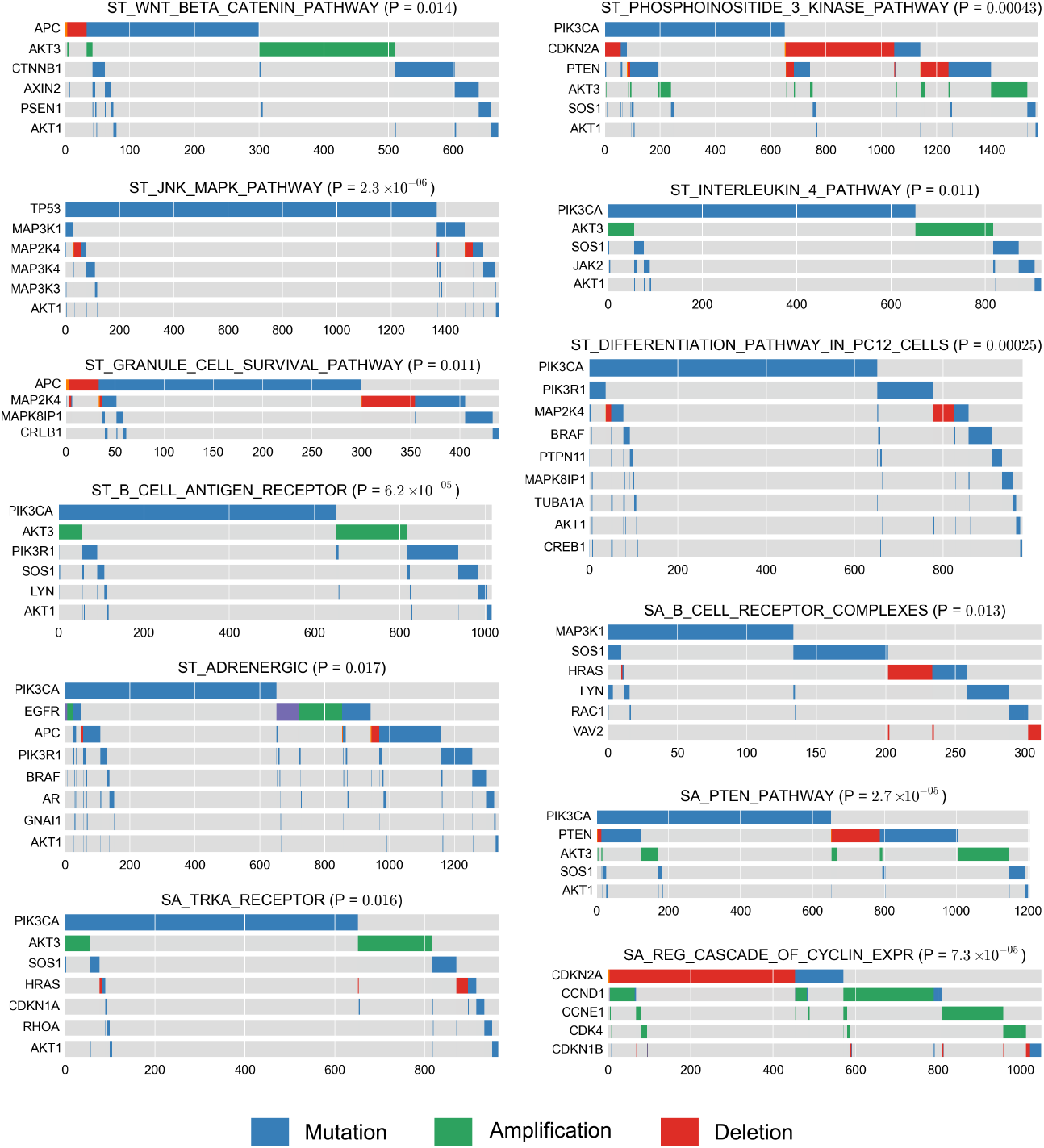
Overview of the significantly mutually exclusive gene sets from the MSigDb canonical pathway collection.

## Supplementary tables

**Supplementary Table 1:**
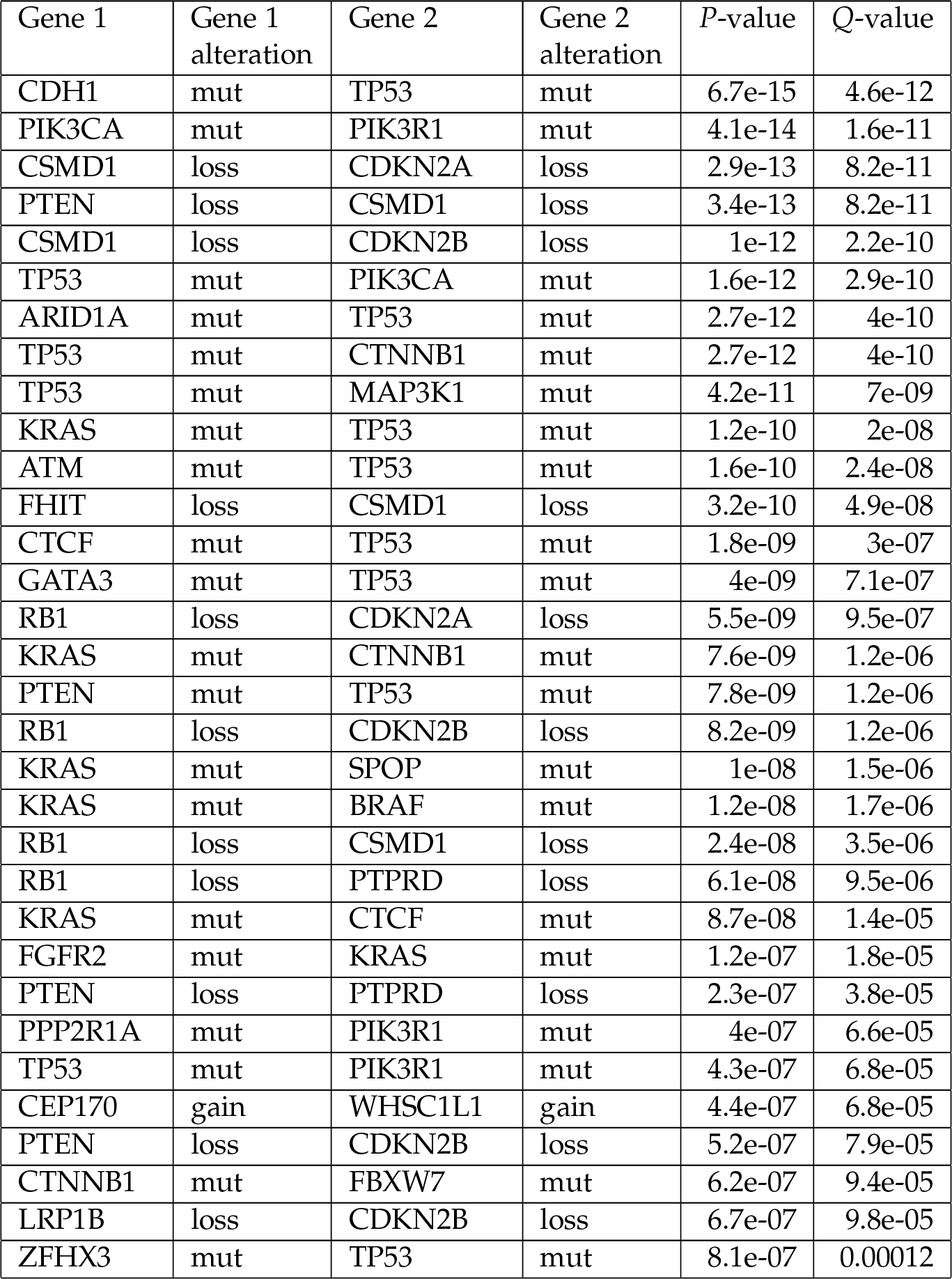
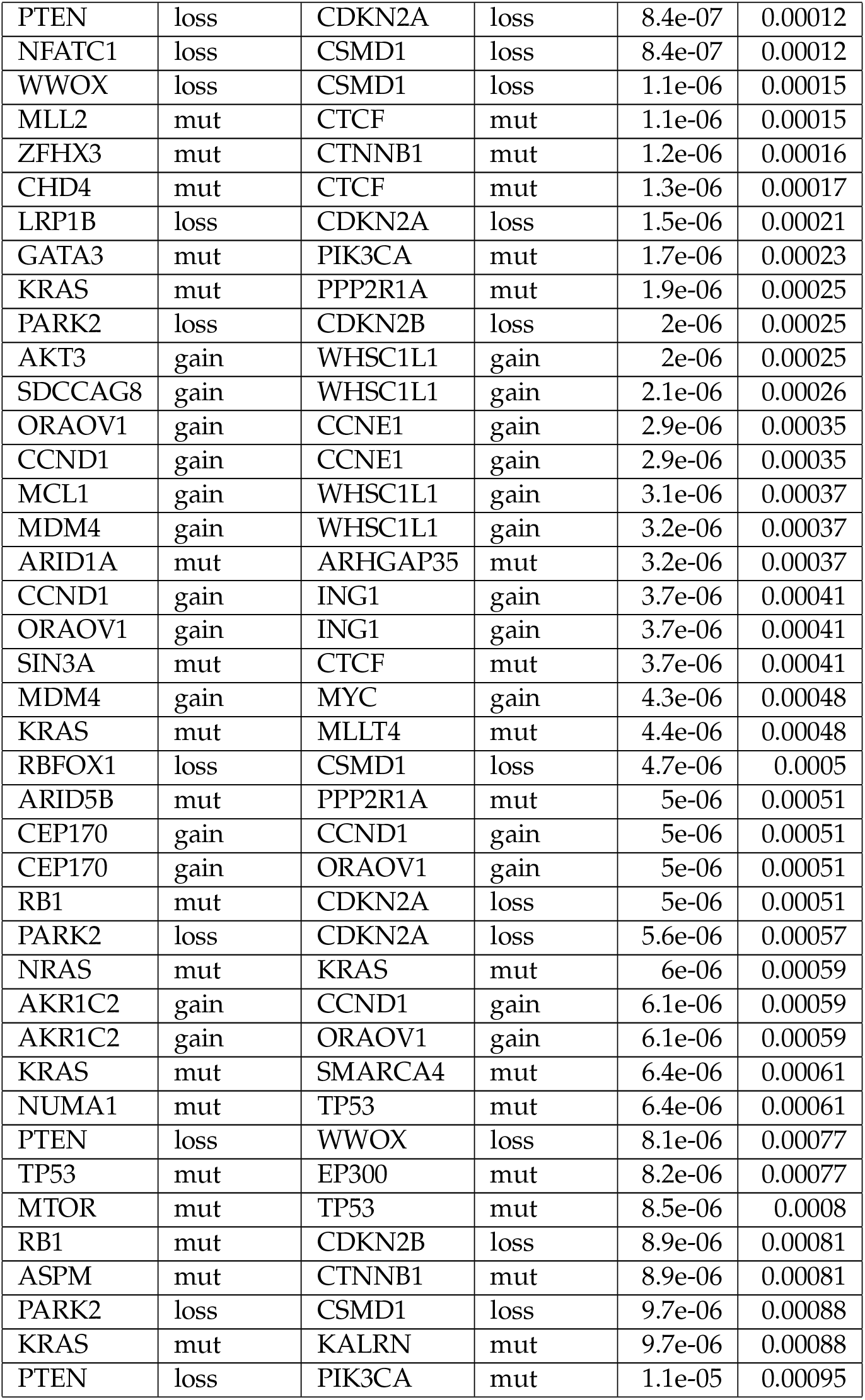
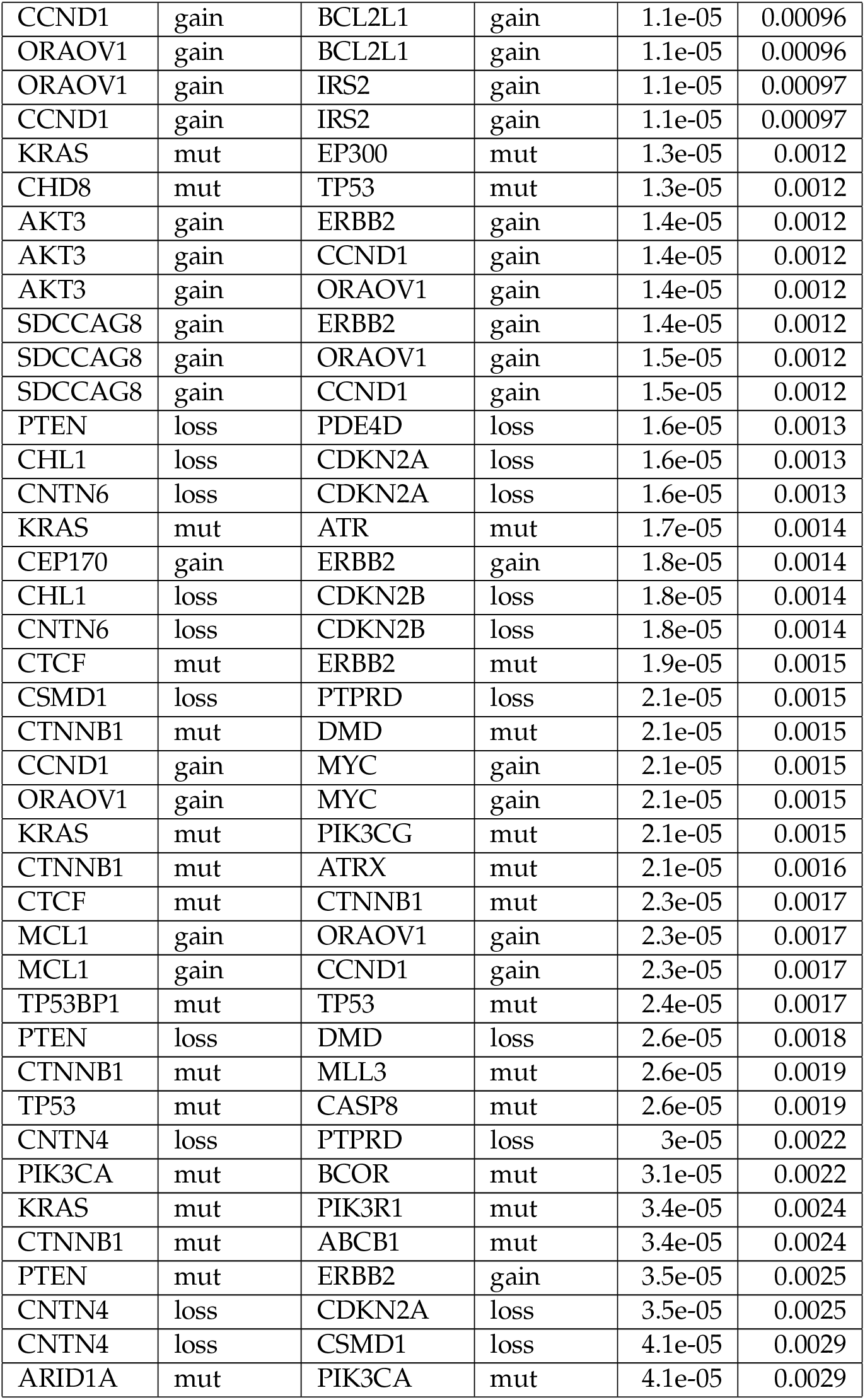
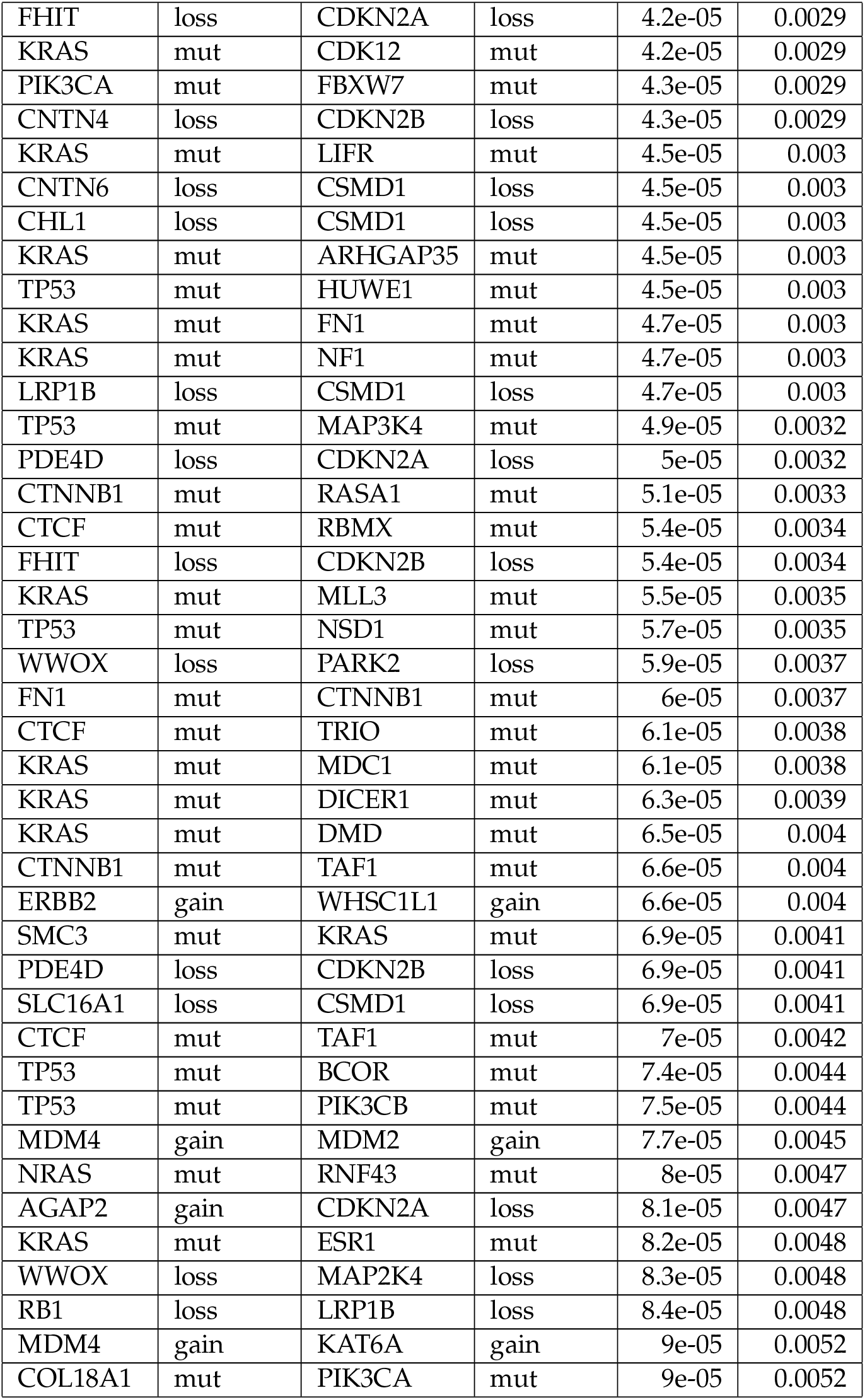
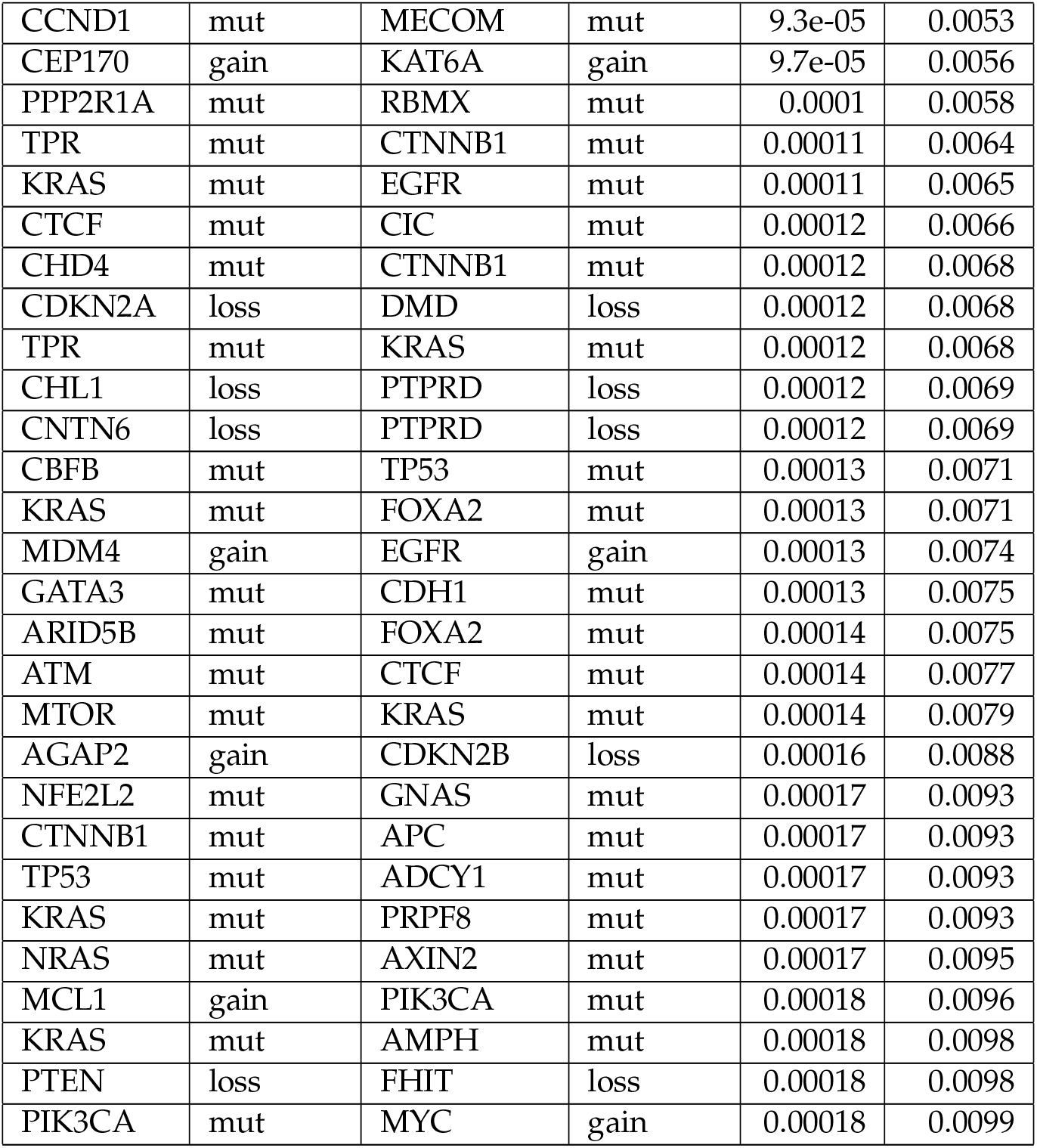
Significantly mutually exclusive alterations found in the pan-cancer data at a maximum FDR of 1%. The first four columns show the genes and the types of alteration involved (mut = point mutation; gain, loss = copy number gain, loss). The *P*-value is obtained using DISCOVER’s mutual exclusivity test. The *Q*-value is the false discover rate estimated using the method of Carlson et al.[44]

| Gene 1 | Gene 1 alteration | Gene 2 | Gene 2 alteration | *P*-value | *Q*-value |
| --- | --- | --- | --- | --- | --- |
| CDH1 | mut | TP53 | mut | 6.7e-15 | 4.6e-12 |
| PIK3CA | mut | PIK3R1 | mut | 4.1e-14 | 1.6e-11 |
| CSMD1 | loss | CDKN2A | loss | 2.9e-13 | 8.2e-11 |
| PTEN | loss | CSMD1 | loss | 3.4e-13 | 8.2e-11 |
| CSMD1 | loss | CDKN2B | loss | 1e-12 | 2.2e-10 |
| TP53 | mut | PIK3CA | mut | 1.6e-12 | 2.9e-10 |
| ARID1A | mut | TP53 | mut | 2.7e-12 | 4e-10 |
| TP53 | mut | CTNNB1 | mut | 2.7e-12 | 4e-10 |
| TP53 | mut | MAP3K1 | mut | 4.2e-11 | 7e-09 |
| KRAS | mut | TP53 | mut | 1.2e-10 | 2e-08 |
| ATM | mut | TP53 | mut | 1.6e-10 | 2.4e-08 |
| FHIT | loss | CSMD1 | loss | 3.2e-10 | 4.9e-08 |
| CTCF | mut | TP53 | mut | 1.8e-09 | 3e-07 |
| GATA3 | mut | TP53 | mut | 4e-09 | 7.1e-07 |
| RB1 | loss | CDKN2A | loss | 5.5e-09 | 9.5e-07 |
| KRAS | mut | CTNNB1 | mut | 7.6e-09 | 1.2e-06 |
| PTEN | mut | TP53 | mut | 7.8e-09 | 1.2e-06 |
| RB1 | loss | CDKN2B | loss | 8.2e-09 | 1.2e-06 |
| KRAS | mut | SPOP | mut | 1e-08 | 1.5e-06 |
| KRAS | mut | BRAF | mut | 1.2e-08 | 1.7e-06 |
| RB1 | loss | CSMD1 | loss | 2.4e-08 | 3.5e-06 |
| RB1 | loss | PTPRD | loss | 6.1e-08 | 9.5e-06 |
| KRAS | mut | CTCF | mut | 8.7e-08 | 1.4e-05 |
| FGFR2 | mut | KRAS | mut | 1.2e-07 | 1.8e-05 |
| PTEN | loss | PTPRD | loss | 2.3e-07 | 3.8e-05 |
| PPP2R1A | mut | PIK3R1 | mut | 4e-07 | 6.6e-05 |
| TP53 | mut | PIK3R1 | mut | 4.3e-07 | 6.8e-05 |
| CEP170 | gain | WHSC1L1 | gain | 4.4e-07 | 6.8e-05 |
| PTEN | loss | CDKN2B | loss | 5.2e-07 | 7.9e-05 |
| CTNNB1 | mut | FBXW7 | mut | 6.2e-07 | 9.4e-05 |
| LRP1B | loss | CDKN2B | loss | 6.7e-07 | 9.8e-05 |
| ZFHX3 | mut | TP53 | mut | 8.1e-07 | 0.00012 |
| PTEN | loss | CDKN2A | loss | 8.4e-07 | 0.00012 |
| NFATC1 | loss | CSMD1 | loss | 8.4e-07 | 0.00012 |
| WWOX | loss | CSMD1 | loss | 1.1e-06 | 0.00015 |
| MLL2 | mut | CTCF | mut | 1.1e-06 | 0.00015 |
| ZFHX3 | mut | CTNNB1 | mut | 1.2e-06 | 0.00016 |
| CHD4 | mut | CTCF | mut | 1.3e-06 | 0.00017 |
| LRP1B | loss | CDKN2A | loss | 1.5e-06 | 0.00021 |
| GATA3 | mut | PIK3CA | mut | 1.7e-06 | 0.00023 |
| KRAS | mut | PPP2R1A | mut | 1.9e-06 | 0.00025 |
| PARK2 | loss | CDKN2B | loss | 2e-06 | 0.00025 |
| AKT3 | gain | WHSC1L1 | gain | 2e-06 | 0.00025 |
| SDCCAG8 | gain | WHSC1L1 | gain | 2.1e-06 | 0.00026 |
| ORAOV1 | gain | CCNE1 | gain | 2.9e-06 | 0.00035 |
| CCND1 | gain | CCNE1 | gain | 2.9e-06 | 0.00035 |
| MCL1 | gain | WHSC1L1 | gain | 3.1e-06 | 0.00037 |
| MDM4 | gain | WHSC1L1 | gain | 3.2e-06 | 0.00037 |
| ARID1A | mut | ARHGAP35 | mut | 3.2e-06 | 0.00037 |
| CCND1 | gain | ING1 | gain | 3.7e-06 | 0.00041 |
| ORAOV1 | gain | ING1 | gain | 3.7e-06 | 0.00041 |
| SIN3A | mut | CTCF | mut | 3.7e-06 | 0.00041 |
| MDM4 | gain | MYC | gain | 4.3e-06 | 0.00048 |
| KRAS | mut | MLLT4 | mut | 4.4e-06 | 0.00048 |
| RBFOX1 | loss | CSMD1 | loss | 4.7e-06 | 0.0005 |
| ARID5B | mut | PPP2R1A | mut | 5e-06 | 0.00051 |
| CEP170 | gain | CCND1 | gain | 5e-06 | 0.00051 |
| CEP170 | gain | ORAOV1 | gain | 5e-06 | 0.00051 |
| RB1 | mut | CDKN2A | loss | 5e-06 | 0.00051 |
| PARK2 | loss | CDKN2A | loss | 5.6e-06 | 0.00057 |
| NRAS | mut | KRAS | mut | 6e-06 | 0.00059 |
| AKR1C2 | gain | CCND1 | gain | 6.1e-06 | 0.00059 |
| AKR1C2 | gain | ORAOV1 | gain | 6.1e-06 | 0.00059 |
| KRAS | mut | SMARCA4 | mut | 6.4e-06 | 0.00061 |
| NUMA1 | mut | TP53 | mut | 6.4e-06 | 0.00061 |
| PTEN | loss | WWOX | loss | 8.1e-06 | 0.00077 |
| TP53 | mut | EP300 | mut | 8.2e-06 | 0.00077 |
| MTOR | mut | TP53 | mut | 8.5e-06 | 0.0008 |
| RB1 | mut | CDKN2B | loss | 8.9e-06 | 0.00081 |
| ASPM | mut | CTNNB1 | mut | 8.9e-06 | 0.00081 |
| PARK2 | loss | CSMD1 | loss | 9.7e-06 | 0.00088 |
| KRAS | mut | KALRN | mut | 9.7e-06 | 0.00088 |
| PTEN | loss | PIK3CA | mut | 1.1e-05 | 0.00095 |
| CCND1 | gain | BCL2L1 | gain | 1.1e-05 | 0.00096 |
| ORAOV1 | gain | BCL2L1 | gain | 1.1e-05 | 0.00096 |
| ORAOV1 | gain | IRS2 | gain | 1.1e-05 | 0.00097 |
| CCND1 | gain | IRS2 | gain | 1.1e-05 | 0.00097 |
| KRAS | mut | EP300 | mut | 1.3e-05 | 0.0012 |
| CHD8 | mut | TP53 | mut | 1.3e-05 | 0.0012 |
| AKT3 | gain | ERBB2 | gain | 1.4e-05 | 0.0012 |
| AKT3 | gain | CCND1 | gain | 1.4e-05 | 0.0012 |
| AKT3 | gain | ORAOV1 | gain | 1.4e-05 | 0.0012 |
| SDCCAG8 | gain | ERBB2 | gain | 1.4e-05 | 0.0012 |
| SDCCAG8 | gain | ORAOV1 | gain | 1.5e-05 | 0.0012 |
| SDCCAG8 | gain | CCND1 | gain | 1.5e-05 | 0.0012 |
| PTEN | loss | PDE4D | loss | 1.6e-05 | 0.0013 |
| CHL1 | loss | CDKN2A | loss | 1.6e-05 | 0.0013 |
| CNTN6 | loss | CDKN2A | loss | 1.6e-05 | 0.0013 |
| KRAS | mut | ATR | mut | 1.7e-05 | 0.0014 |
| CEP170 | gain | ERBB2 | gain | 1.8e-05 | 0.0014 |
| CHL1 | loss | CDKN2B | loss | 1.8e-05 | 0.0014 |
| CNTN6 | loss | CDKN2B | loss | 1.8e-05 | 0.0014 |
| CTCF | mut | ERBB2 | mut | 1.9e-05 | 0.0015 |
| CSMD1 | loss | PTPRD | loss | 2.1e-05 | 0.0015 |
| CTNNB1 | mut | DMD | mut | 2.1e-05 | 0.0015 |
| CCND1 | gain | MYC | gain | 2.1e-05 | 0.0015 |
| ORAOV1 | gain | MYC | gain | 2.1e-05 | 0.0015 |
| KRAS | mut | PIK3CG | mut | 2.1e-05 | 0.0015 |
| CTNNB1 | mut | ATRX | mut | 2.1e-05 | 0.0016 |
| CTCF | mut | CTNNB1 | mut | 2.3e-05 | 0.0017 |
| MCL1 | gain | ORAOV1 | gain | 2.3e-05 | 0.0017 |
| MCL1 | gain | CCND1 | gain | 2.3e-05 | 0.0017 |
| TP53BP1 | mut | TP53 | mut | 2.4e-05 | 0.0017 |
| PTEN | loss | DMD | loss | 2.6e-05 | 0.0018 |
| CTNNB1 | mut | MLL3 | mut | 2.6e-05 | 0.0019 |
| TP53 | mut | CASP8 | mut | 2.6e-05 | 0.0019 |
| CNTN4 | loss | PTPRD | loss | 3e-05 | 0.0022 |
| PIK3CA | mut | BCOR | mut | 3.1e-05 | 0.0022 |
| KRAS | mut | PIK3R1 | mut | 3.4e-05 | 0.0024 |
| CTNNB1 | mut | ABCB1 | mut | 3.4e-05 | 0.0024 |
| PTEN | mut | ERBB2 | gain | 3.5e-05 | 0.0025 |
| CNTN4 | loss | CDKN2A | loss | 3.5e-05 | 0.0025 |
| CNTN4 | loss | CSMD1 | loss | 4.1e-05 | 0.0029 |
| ARID1A | mut | PIK3CA | mut | 4.1e-05 | 0.0029 |
| FHIT | loss | CDKN2A | loss | 4.2e-05 | 0.0029 |
| KRAS | mut | CDK12 | mut | 4.2e-05 | 0.0029 |
| PIK3CA | mut | FBXW7 | mut | 4.3e-05 | 0.0029 |
| CNTN4 | loss | CDKN2B | loss | 4.3e-05 | 0.0029 |
| KRAS | mut | LIFR | mut | 4.5e-05 | 0.003 |
| CNTN6 | loss | CSMD1 | loss | 4.5e-05 | 0.003 |
| CHL1 | loss | CSMD1 | loss | 4.5e-05 | 0.003 |
| KRAS | mut | ARHGAP35 | mut | 4.5e-05 | 0.003 |
| TP53 | mut | HUWE1 | mut | 4.5e-05 | 0.003 |
| KRAS | mut | FN1 | mut | 4.7e-05 | 0.003 |
| KRAS | mut | NF1 | mut | 4.7e-05 | 0.003 |
| LRP1B | loss | CSMD1 | loss | 4.7e-05 | 0.003 |
| TP53 | mut | MAP3K4 | mut | 4.9e-05 | 0.0032 |
| PDE4D | loss | CDKN2A | loss | 5e-05 | 0.0032 |
| CTNNB1 | mut | RASA1 | mut | 5.1e-05 | 0.0033 |
| CTCF | mut | RBMX | mut | 5.4e-05 | 0.0034 |
| FHIT | loss | CDKN2B | loss | 5.4e-05 | 0.0034 |
| KRAS | mut | MLL3 | mut | 5.5e-05 | 0.0035 |
| TP53 | mut | NSD1 | mut | 5.7e-05 | 0.0035 |
| WWOX | loss | PARK2 | loss | 5.9e-05 | 0.0037 |
| FN1 | mut | CTNNB1 | mut | 6e-05 | 0.0037 |
| CTCF | mut | TRIO | mut | 6.1e-05 | 0.0038 |
| KRAS | mut | MDC1 | mut | 6.1e-05 | 0.0038 |
| KRAS | mut | DICER1 | mut | 6.3e-05 | 0.0039 |
| KRAS | mut | DMD | mut | 6.5e-05 | 0.004 |
| CTNNB1 | mut | TAF1 | mut | 6.6e-05 | 0.004 |
| ERBB2 | gain | WHSC1L1 | gain | 6.6e-05 | 0.004 |
| SMC3 | mut | KRAS | mut | 6.9e-05 | 0.0041 |
| PDE4D | loss | CDKN2B | loss | 6.9e-05 | 0.0041 |
| SLC16A1 | loss | CSMD1 | loss | 6.9e-05 | 0.0041 |
| CTCF | mut | TAF1 | mut | 7e-05 | 0.0042 |
| TP53 | mut | BCOR | mut | 7.4e-05 | 0.0044 |
| TP53 | mut | PIK3CB | mut | 7.5e-05 | 0.0044 |
| MDM4 | gain | MDM2 | gain | 7.7e-05 | 0.0045 |
| NRAS | mut | RNF43 | mut | 8e-05 | 0.0047 |
| AGAP2 | gain | CDKN2A | loss | 8.1e-05 | 0.0047 |
| KRAS | mut | ESR1 | mut | 8.2e-05 | 0.0048 |
| WWOX | loss | MAP2K4 | loss | 8.3e-05 | 0.0048 |
| RB1 | loss | LRP1B | loss | 8.4e-05 | 0.0048 |
| MDM4 | gain | KAT6A | gain | 9e-05 | 0.0052 |
| COL18A1 | mut | PIK3CA | mut | 9e-05 | 0.0052 |
| CCND1 | mut | MECOM | mut | 9.3e-05 | 0.0053 |
| CEP170 | gain | KAT6A | gain | 9.7e-05 | 0.0056 |
| PPP2R1A | mut | RBMX | mut | 0.0001 | 0.0058 |
| TPR | mut | CTNNB1 | mut | 0.00011 | 0.0064 |
| KRAS | mut | EGFR | mut | 0.00011 | 0.0065 |
| CTCF | mut | CIC | mut | 0.00012 | 0.0066 |
| CHD4 | mut | CTNNB1 | mut | 0.00012 | 0.0068 |
| CDKN2A | loss | DMD | loss | 0.00012 | 0.0068 |
| TPR | mut | KRAS | mut | 0.00012 | 0.0068 |
| CHL1 | loss | PTPRD | loss | 0.00012 | 0.0069 |
| CNTN6 | loss | PTPRD | loss | 0.00012 | 0.0069 |
| CBFB | mut | TP53 | mut | 0.00013 | 0.0071 |
| KRAS | mut | FOXA2 | mut | 0.00013 | 0.0071 |
| MDM4 | gain | EGFR | gain | 0.00013 | 0.0074 |
| GATA3 | mut | CDH1 | mut | 0.00013 | 0.0075 |
| ARID5B | mut | FOXA2 | mut | 0.00014 | 0.0075 |
| ATM | mut | CTCF | mut | 0.00014 | 0.0077 |
| MTOR | mut | KRAS | mut | 0.00014 | 0.0079 |
| AGAP2 | gain | CDKN2B | loss | 0.00016 | 0.0088 |
| NFE2L2 | mut | GNAS | mut | 0.00017 | 0.0093 |
| CTNNB1 | mut | APC | mut | 0.00017 | 0.0093 |
| TP53 | mut | ADCY1 | mut | 0.00017 | 0.0093 |
| KRAS | mut | PRPF8 | mut | 0.00017 | 0.0093 |
| NRAS | mut | AXIN2 | mut | 0.00017 | 0.0095 |
| MCL1 | gain | PIK3CA | mut | 0.00018 | 0.0096 |
| KRAS | mut | AMPH | mut | 0.00018 | 0.0098 |
| PTEN | loss | FHIT | loss | 0.00018 | 0.0098 |
| PIK3CA | mut | MYC | gain | 0.00018 | 0.0099 |

## Supplementary text

### Parameter estimation

To apply the DISCOVER test, we need estimates of the alteration probabilities *p_ij_* for all genes *i* and all tumors *j*. These estimates are used as parameters of the Poisson-Binomial distribution used by the test. Here, we show that each alteration probability can be written in terms of two parameters.

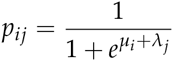

For estimating the parameters of the Poisson-Binomial distribution we maximize the information entropy (or equivalently below, minimize the negative of the entropy), subject to the constraints that the expected row and column marginals match the observed ones.

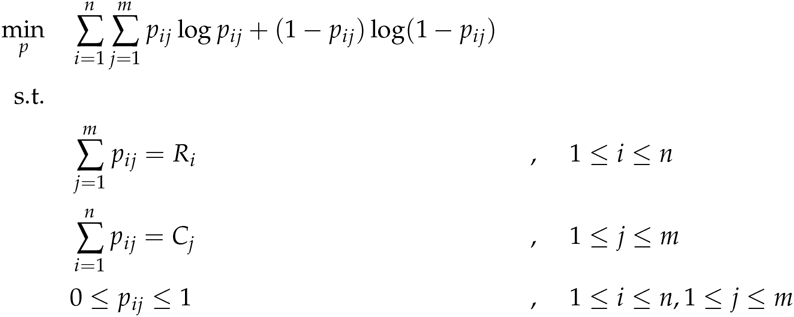

where

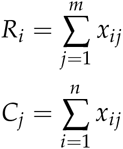

We first turn this constrained optimization problem into an unconstrained one by defining the Lagrangian dual. The Lagrangian is as follows.

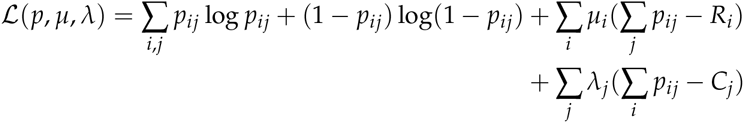

which we rewrite to

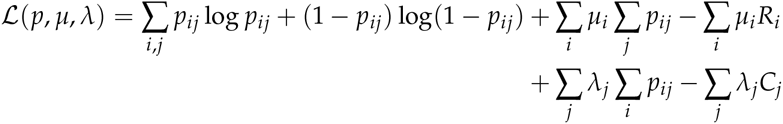

We then optimize the Lagrangian with respect to the variables *p_ij_* by setting the partial derivatives to 0 and solving for those variables.

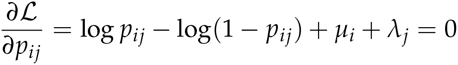

From this, we derive that

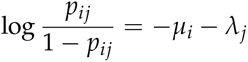

The left-hand side of this equation is the familiar logit function, the inverse of which is the logistic function. Hence, we obtain the following expression for *p_ij_*.

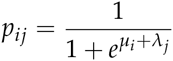

With this, we can formulate the Lagrangian dual as follows.

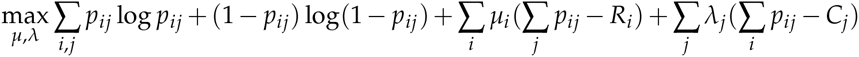

where *p_ij_* is defined as above.

